# Prioritization of autoimmune disease-associated genetic variants that perturb regulatory element activity in T cells

**DOI:** 10.1101/2021.05.30.445673

**Authors:** Kousuke Mouri, Michael H. Guo, Carl G. de Boer, Gregory A. Newby, Matteo Gentili, David R. Liu, Nir Hacohen, Ryan Tewhey, John P. Ray

**Affiliations:** The Jackson Laboratory, Bar Harbor, ME 04609; Department of Neurology, Perelman School of Medicine, University of Pennsylvania, Philadelphia, PA 19104; Broad Institute of Harvard and MIT, Cambridge, MA 02142; School of Biomedical Engineering, University of British Columbia, Vancouver, BC, Canada V6T 1Z3; Department of Chemistry and Chemical Biology, Harvard University, Cambridge, MA 02138; Howard Hughes Medical Institute, Harvard University, Cambridge, MA 02138; Center for Cancer Research, Massachusetts General Hospital, Boston, MA, 02114; Systems Immunology, Benaroya Research Institute, Seattle, WA 98101

## Abstract

Genome-wide association studies have uncovered hundreds of autoimmune disease-associated loci; however, the causal genetic variant(s) within each locus are mostly unknown. Here, we perform high-throughput allele-specific reporter assays to prioritize disease-associated variants for five autoimmune diseases. By examining variants that both promote allele-specific reporter expression and are located in accessible chromatin, we identify 60 putatively causal variants that enrich for statistically fine-mapped variants by up to 57.8-fold. We introduced the risk allele of a prioritized variant (rs72928038) into a human T cell line and deleted the orthologous sequence in mice, both resulting in reduced *BACH2* expression. Naïve CD8 T cells from mice containing the deletion had reduced expression of genes that suppress activation and maintain stemness. Our results represent an example of an effective approach for prioritizing variants and studying their physiologically relevant effects.

## INTRODUCTION

Genome-wide association studies (GWAS) are a powerful approach for identifying genetic susceptibility loci for autoimmune diseases. However, our ability to draw direct mechanistic insights from GWAS loci has been hampered by challenges in identifying which variant(s) actually cause disease risk at any given locus. Pinpointing the specific causal variant provides insight into the context and mechanism by which the disease association modulates disease risk. There are three major challenges to identifying causal variant(s): 1) at most loci, there are many disease-associated variants due to linkage disequilibrium (LD) between causal and non-causal variants, 2) ∼90% of causal variants reside in non-coding regions^1,2^, where their mechanisms of action are difficult to infer, and 3) the context (e.g., cell-type, cell-state, etc.) in which variants act may at times be difficult to discern, particularly for non-coding variants. Thus, to discern causal variants, we must refine strategies to prioritize and test variants for how they perturb genomic functions, particularly in disease-relevant cell types and states.

Recent methodologies have been developed to distinguish causal variants from those that are non-causal, including inferring the cell types in which they act. Statistical fine-mapping methods can generate credible sets of likely causal variants, with high powered studies able to pinpoint singular causal variants for many disease associations^3^; however, most disease association studies lack sufficient power to definitively distinguish the causal variant(s) for each locus. Experimental and computational methodologies have also been developed to discern putatively causal variants and infer the context in which they act. For instance, overlaying maps of accessible and active chromatin regions through DNase Hypersensitivity I and H3K27ac ChIP-sequencing in many cell types and environmental conditions have enriched for likely causal variants, and these methods can aid in identifying disease-relevant cell types^1,2,4,5^. In addition, perturbational studies-- such as massively parallel reporter assays (MPRA) that test genetic variants for their ability to modulate gene expression and CRISPR inhibition which perturbs putative regulatory elements to determine their effect on gene expression-- also enrich for disease-associated variants^6,7^. While each of these methodologies is useful for prioritizing potential causal variants, they all have imperfect accuracy due to differences in how variants act in the context of the assay as compared to disease-relevant states^6^. Thus, once variants are prioritized, they require further mechanistic dissection to determine whether they are causal for disease associations, such as through editing the variants into the genomes of disease-relevant cells.

Here, we used a highly efficacious prioritization scheme on ∼18,000 variants associated with five autoimmune diseases including type 1 diabetes (T1D), inflammatory bowel disease ([IBD], including ulcerative colitis [UC] and Crohn’s disease [CD]), rheumatoid arthritis (RA), psoriasis, and multiple sclerosis (MS) to identify likely causal variants. Through integrating MPRA and chromatin accessibility data, we found 60 likely causal variants that enriched up to 57.8-fold for causal variants according to fine-mapping. We further characterized the effects of a single variant (rs72928038) associated with multiple autoimmune diseases through analyzing the presence of the risk allele in accessible chromatin, using a base editing approach to insert the variant into a human T cell line, and by constructing mice containing a deletion at the orthologous genomic region. Human T cells heterozygous for the variant have substantial reductions in accessible chromatin containing the risk allele, and insertion of the variant into Jurkat T cells reduced expression of *BACH2*, a transcriptional repressor that negatively regulates effector T cell differentiation^8^, and positively regulates regulatory T cell differentiation^9^ and T cell stemness^10^. Because the region containing rs72928038 is highly conserved, and the orthologous region in mouse is also in a putative cis-regulatory element in mouse T cells, we created mice with a small deletion in the non-coding region overlapping rs72928038 to determine its effect on *Bach2* and global expression of naïve T cells. We found rs72928038-deleted mice to have naïve CD8 T cells with reduced *Bach2* expression and reduced expression of naïve T cell stemness-associated genes, indicating that rs72928038 plays an important role in suppressing naïve T cell activation. This work demonstrates a framework for combining chromatin accessibility and MPRA to identify variants that impact risk for autoimmune disease and provides a clear example of how to move from variant prioritization to causal effects on cellular outcome in an organismic model.

## RESULTS

### Prioritizing autoimmune GWAS variants with MPRA

Because autoimmune disease-associated genetic variants are highly enriched in T cell cis-regulatory elements^1,2,5,6,11,12^, we hypothesized that many disease-causal variants likely alter the activity of T cell cis-regulatory elements. One way to test the effect of variants on regulatory activities is through testing variant alleles for their differential effects on reporter expression in MPRA^13–16^. To this end, we created MPRA libraries for variants associated with diseases in which T cells are known to play a role (henceforth collectively referred to as T-GWAS). These diseases include IBD (including CD and UC)^17^, MS^18,19^, T1D^20^, psoriasis^21^, and RA^22^.

We collected 578 GWAS index variants (representing 531 distinct GWAS loci) and variants in tight LD (r^2^ > 0.8) from the above-cited studies, totaling 18,312 variants (Supplementary Table 1). To generate our MPRA, library alleles were synthesized as 200 bp elements centered within their genomic context. We also included 91 positive enhancer controls and 506 negative controls used in a previous MPRA study (Supplementary Table 2)^13^. We barcoded the enhancer elements (∼1,000 barcodes/element) and cloned them into the MPRA vector, followed by inserting a minimal promoter and GFP (Fig. 1a, Supplementary Fig.1a, Methods). After nucleofection of the library into Jurkat T cells, followed by RNA-sequencing of barcodes after 24 h, we found that barcode prevalence in plasmid and cDNA replicates was tightly correlated, and that some barcodes were more present in cDNA than in plasmid libraries, indicative of their higher expression (Supplementary Fig.1b; Supplementary Table 3). We found 7,095 elements that had higher reporter expression than expected from their prevalence in plasmid libraries for at least one variant allele (termed putative cis-regulatory elements, pCREs; Supplementary Fig.1c; Supplementary Table 4); positive enhancer controls generally were pCREs, while negative controls had minimal expression (Supplementary Fig.1d). Of the 7,095 pCRE elements, we found 313 variants that had statistically significant differences in expression between the reference and alternate alleles, which we term expression-modulating variants (emVars) (Fig. 1b; Supplementary Table 4).

**Fig. 1.**
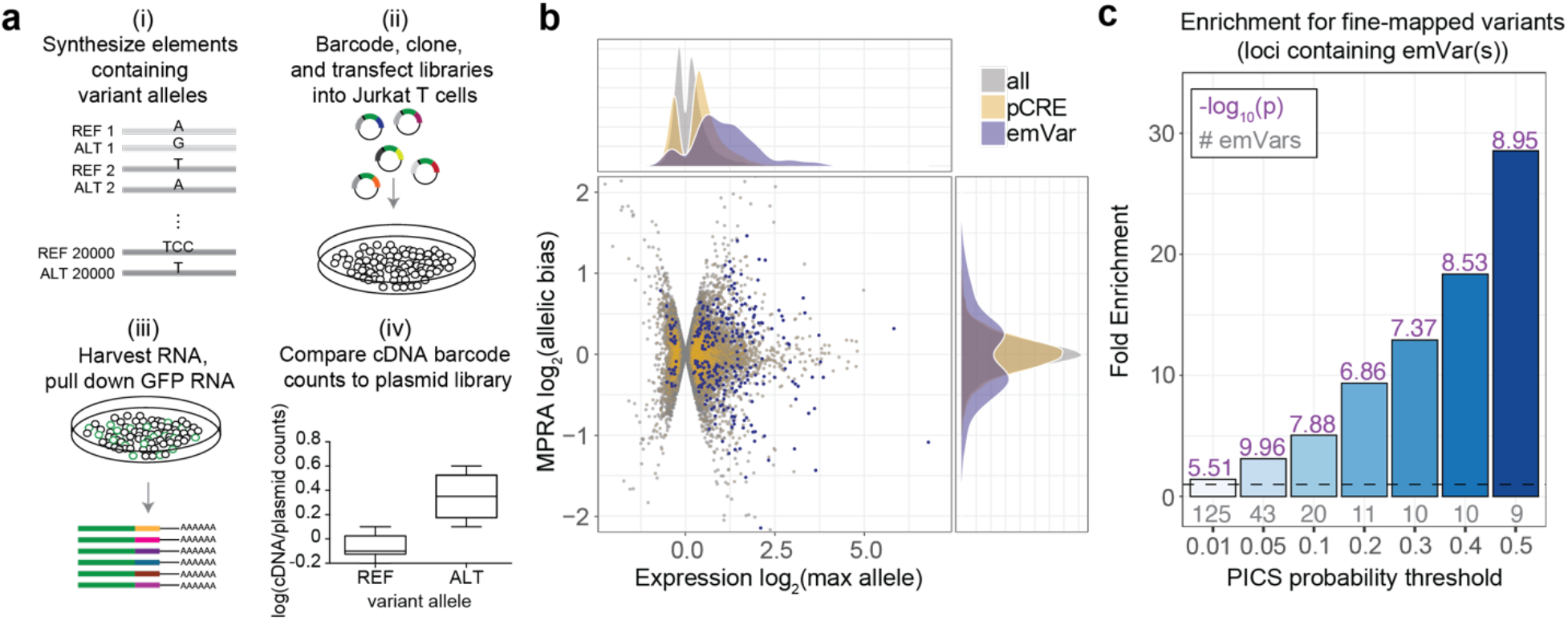
Prioritizing GWAS variants using high-throughput reporter assays in Jurkat T cells. **a**) Workflow for creating MPRA libraries-i) Oligonucleotide synthesis of variants and 200 bp surrounding genomic region; ii) barcoding, cloning, and transfection of plasmid library into Jurkat T cells; iii) harvesting RNA from Jurkat and pull down of GFP mRNA; iv) RNA-sequencing of barcodes, normalization to their prevalence in the plasmid library, and comparison of alleles for differential reporter activity (a more detailed workflow is provided in Supplementary Fig. 1a). **b**) Volcano plot. The log_2_ expression value of the highest expressing allele is on the X axis, and the log_2_ of the activity of allele1/allele2 is on the Y axis. pCRE = putative cis-regulatory element; emVar = expression-modulating variant. **c**) Bar plot showing enrichment of emVars for PICS statistically fine-mapped variants at GWAS loci where an emVar was detected, with the minimum PICS probability threshold indicated on the X axis. Gray numbers below each bar show the number of emVars that are statistically fine-mapped at a given PICS probability threshold. Purple numbers above each bar show the -log_10_ of the enrichment P value. Details of PICS enrichment results are shown in Table S9. Enrichment in (**c**) was calculated as a risk ratio (see Methods), and P values were determined through a two-sided Fisher’s exact test.

We next assessed whether there are specific cis-regulatory phenotypes in which emVars were preferentially found. We observed emVars were most highly enriched in transcription start site (TSS) regions and distal enhancers (Supplementary Fig. 2a and b), with particularly high enrichment at regions marked by H3K4me3, CAGE and DNase hypersensitivity sites (DHS) (Supplementary Fig. 2c), consistent with many emVars altering regulatory element activity. emVars were also more likely to have allelic bias in ATAC-seq data from hematopoietic cell types and to be a chromatin accessibility quantitative trait locus (caQTL) as compared to MPRA variants with no activity, and emVar allelic effects were correlated with allelic bias and QTL directionality from these data (Supplementary Fig. 3). Consistent with emVars disrupting regulatory element activity and chromatin accessibility, we found that their allelic effects were correlated with computationally predicted allelic effects (“delta SVM”) in CD4 T cell enhancer elements, with most emVars showing directional concordance with the delta SVM score (Supplementary Fig. 4a). emVars were also much more likely to perturb a transcription factor (TF) motif (according to position weight matrices) when compared to all variants tested in the MPRA assay, with predicted TF binding also correlating strongly with the observed MPRA allelic bias for emVars (Supplementary Fig. 4b-d; Supplementary Tables 4-6). Therefore, T-GWAS emVars enrich in regulatory regions and for variants that have orthogonal regulatory phenotypes and allele-specific activities.

Since emVars enrich for variants that impact regulatory activity, we predicted that MPRA could be used to identify causal variants at GWAS loci. Most GWAS loci are thought to have one or a small number of causal variants, with remaining variants statistically associated with a given disease solely due to tight LD with the true causal variant(s). Consistent with this notion, of the 181 GWAS loci for which we found an emVar (31% of all assessed GWAS loci; Supplementary Fig. 5a), 120/181 loci had only one emVar, and 169/181 loci had four or fewer emVars (Supplementary Fig. 5b).

To test if emVars are identifying causal variants, we next tested whether emVars are enriched for variants identified by statistical fine-mapping. We performed fine-mapping using PICS^1^ for all five autoimmune diseases (Supplementary Tables 7 and 8). Among the various statistical fine-mapping approaches available, we chose to use PICS as it does not require full GWAS summary statistics, which were unavailable for many of the diseases we analyzed. We tested whether emVars are enriched for high posterior probability variants at various posterior probability thresholds. When taking into account all GWAS loci, regardless of whether an emVar was identified, emVars enriched up to 3.49-fold for causal variants according to PICS (Supplementary Fig. 6a; Supplementary Table 9). With an understanding that MPRA will not identify variants in all loci such as those where a coding variant is causal, we decided to test enrichments for loci where at least one emVar was identified. Within loci where we identified at least one emVar, we found that emVars were as much as 28.5-fold enriched for causal variants according to PICS (Fig. 1c; Supplementary Table 10). Among loci containing both an emVar and a high posterior probability fine-mapped SNP (posterior inclusion probability [PIP] > 0.5), 45% of the high PIP fine-mapped SNPs were also emVars. Since the T1D GWAS we used to create our MPRA library also contained statistical fine-mapping data^20^, we assessed the enrichment of emVars for statistically fine-mapped variants from this separate dataset, detecting up to a 4.17-fold enrichment (Supplementary Fig. 6b). These data suggest that MPRA is a highly robust approach for prioritizing causal disease variants.

To estimate the sensitivity and specificity of the MPRA, we again leveraged PICS statistical fine-mapping. We constructed credible sets of statistically fine-mapped variants (for example, an 80% credible set will contain the causal variant 80% of the time (Supplementary Table 7; see Methods)), similar to the approach utilized in a recent MPRA study^14^. At various PICS credible sets, we calculated the sensitivity of the MPRA to be 18.4% to 19.7.%, and specificity ranging from 90.8% to 95.0%. Thus, MPRA can prioritize causal variants at a fifth of all loci while maintaining high specificity.

### emVars in T cell accessible chromatin occur near genes that regulate T cell function

We found that active elements within our MPRA were enriched for regions of accessible chromatin from T cells and other hematopoietic cell types compared to non-hematopoietic cell types (Fig. 2a), suggesting that MPRA regulatory activity accurately reflects the transcriptional regulation of the cell type in which it is tested. However, because MPRA evaluates the regulatory activity of a sequence outside its genomic context, effects of variants measured by MPRA may differ from the true endogenous effects. We previously performed MPRA on all common variants near *TNFAIP3* and observed a higher enrichment for putatively disease-causal variants when taking the intersection of MPRA results and regions of accessible chromatin^6^. To assess whether these prior findings extend to our genome-wide T-GWAS MPRA experiment, we compared all variants tested in MPRA to those that were in DHS regions in T cells. Of the 313 emVars, 60 overlapped a T cell DHS peak (Supplementary Table 4). For genetic associations that had at least one emVar in accessible chromatin, we found up to 57.8-fold enrichment for causal variants according to PICS (Fig. 2b and Supplementary Table 10; when taking into account all GWAS loci tested, there was a 9.3-fold enrichment for causal variants according to PICS, Supplementary Fig. 7 and Supplementary Table 9). We calculated sensitivity and specificity for emVars within PICS credible sets at loci where any variant on the haplotype overlapped a T cell DHS peak. When subsetting loci for those with a variant in DHS, MPRA achieved a sensitivity ranging from 23.1% to 25.5% and specificity ranging from 81.1% to 91.7%. Therefore, emVars that are present in the accessible chromatin of T cells enriched strongly for causal variants, and to a much greater extent than either methodology alone.

**Fig. 2.**
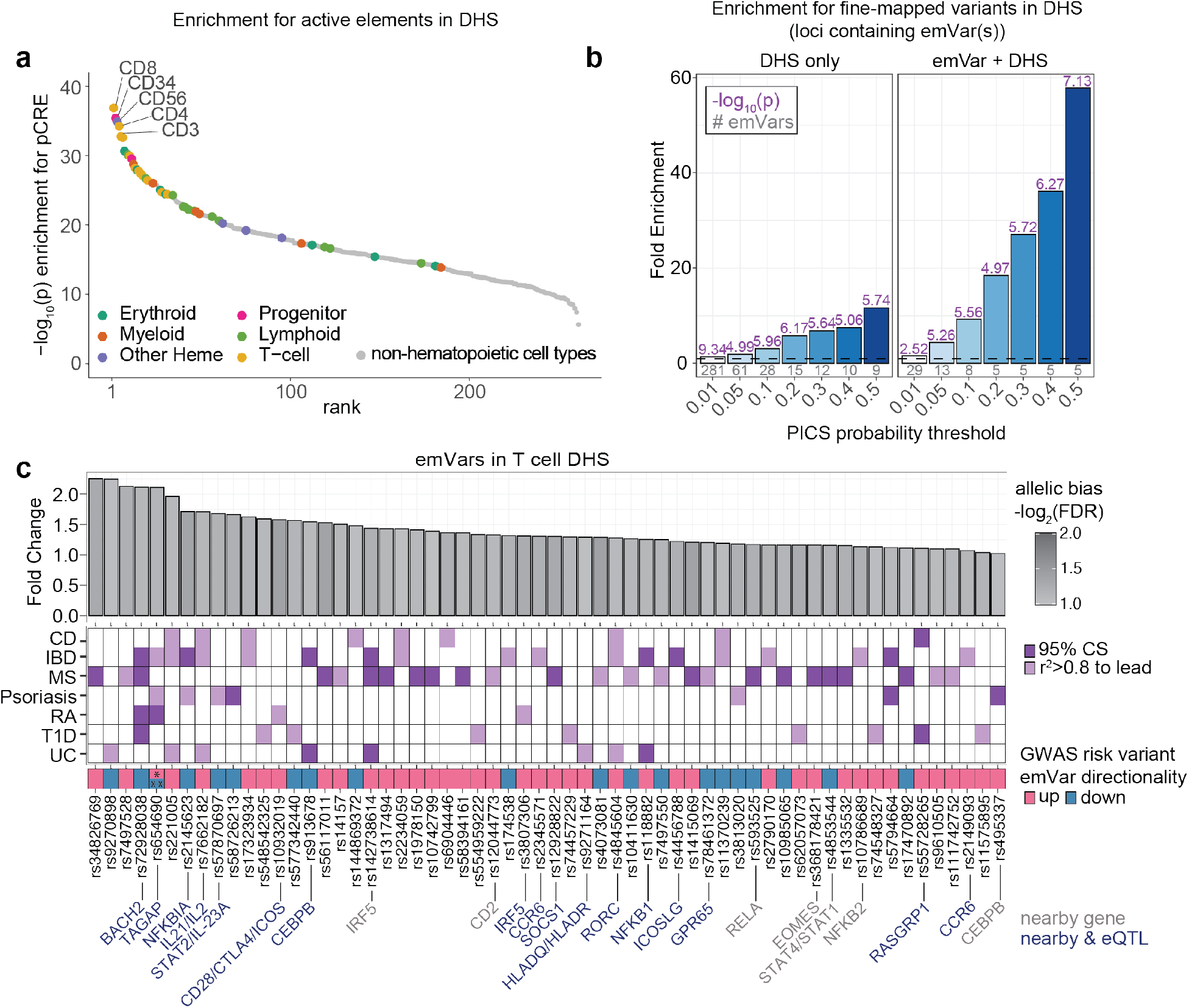
T-GWAS emVars in T cell accessible chromatin enrich highly for fine-mapped variants. **a**) Enrichment of DHS sites from hematopoietic and non-hematopoietic cell types for MPRA pCREs. Cell types are ranked from left to right from most statistically significant to least significant. Hematopoietic cell types are colored by their ontogeny as indicated in the legend. Non-hematopoietic cell types are shown in gray. Y-axis shows the -log_10_ of the enrichment P - value. **b**) Enrichment of statistically fine-mapped variants within T cell DHS sites (left) and enrichment of statistically fine-mapped variants that are emVars within T cell DHS sites (right). Details of PICS enrichment results are shown in Table S9. **c**) Bar plot (top) of 60 emVars in T cell DHS sites with their allelic bias (y-axis) and log_2_FDR (shade of bar). GWAS for which emVar is associated (middle). emVars in 95% fine-mapping credible sets are shown in dark purple, while variants in tight LD to lead the variant (r^2^ > 0.8) but not in credible sets are shown in light purple. Immediately underneath, pink and teal boxes indicate the MPRA expression directionality of the GWAS disease risk-increasing variant as compared to the non-risk variant, followed by variant rsIDs. For one variant, rs654690, the risk alleles are opposing depending on disease, with * indicating the risk allele for both psoriasis and IBD, and ** indicating the risk allele for RA. Nearby genes that are known to play a role in T cell differentiation and function (gray) and nearby genes for which the variant is an eQTL (dark blue; according to Open Targets Genetics;^55^ are listed on bottom. Enrichment (**a**) were determined through a two-sided Fisher’s exact test. Enrichment in (**b**) was calculated as a risk ratio (see Methods), and P values were determined through a two-sided Fisher’s exact test. Statistical significance of allelic bias in (**c, top bar plot**) was calculated using a paired Student’s two-sided *t*-test.

Many emVars in accessible chromatin were near (and in most cases were eQTLs for) genes with important roles in T cell biology, including genes that regulate T cell differentiation (*BACH2, EOMES, RORC, CEBPB*), signal transduction (*CD28, CTLA4, ICOS, STAT1, STAT2, STAT4, IRF5, NFKB1, NFKB2, RELA, SOCS1*), cytokine production (*IL2, IL21, IL23*), and migration (*CCR6*) (Fig. 2c). rs654690, associated with psoriasis, IBD, and RA, falls preferentially in the accessible chromatin of Tregs and contacts the *TAGAP* promoter ∼50kb downstream^23^; *TAGAP* is a gene that has been shown to play a role in Th17 cell differentiation and thymocyte trafficking^24–26^ (Fig. 3a). The rs654690 disease-risk allele depends on the disease: the RA risk-increasing allele (C) decreases MPRA activity, while the psoriasis and IBD risk-increasing allele (T) increases MPRA activity (Fig. 2c). Disease-risk alleles for two variants, rs142738614, associated with MS, RA, and UC, and rs3807306, associated with RA, were in moderate LD to each other (*r*^2^ = 0.7), and in separate regulatory elements of *IRF5*, a gene with many roles in immunity, including T cell-intrinsic roles that modulate signaling, migration, and differentiation^27^ (Supplementary Fig. 8a). The risk alleles for both variants drove an increase in reporter expression (Fig. 2c). rs55728265, associated with CD and T1D, is in the 5′ UTR of *RASGRP1*, a gene that regulates T cell signaling and differentiation^28,29^ (Supplementary Fig. 8b); the rs55728265 risk allele increases reporter expression (Fig. 2c). rs72928038, associated with T1D, RA, and MS, is within an intron of *BACH2*, a gene involved in suppressing effector CD4 and CD8 T cell differentiation, while promoting regulatory T cell differentiation^8,9^ and T cell stemness^10^ (Fig. 3b). This variant falls preferentially within the accessible chromatin of naïve T cells and contacts the *BACH2* promoter in naïve T cells^23^, with the risk allele reducing reporter expression (Fig. 2c). Collectively, these data suggest that disease-associated emVars that act in T cell regulatory regions regulate genes known to play a role in T cell signaling, differentiation, and function.

**Fig. 3.**
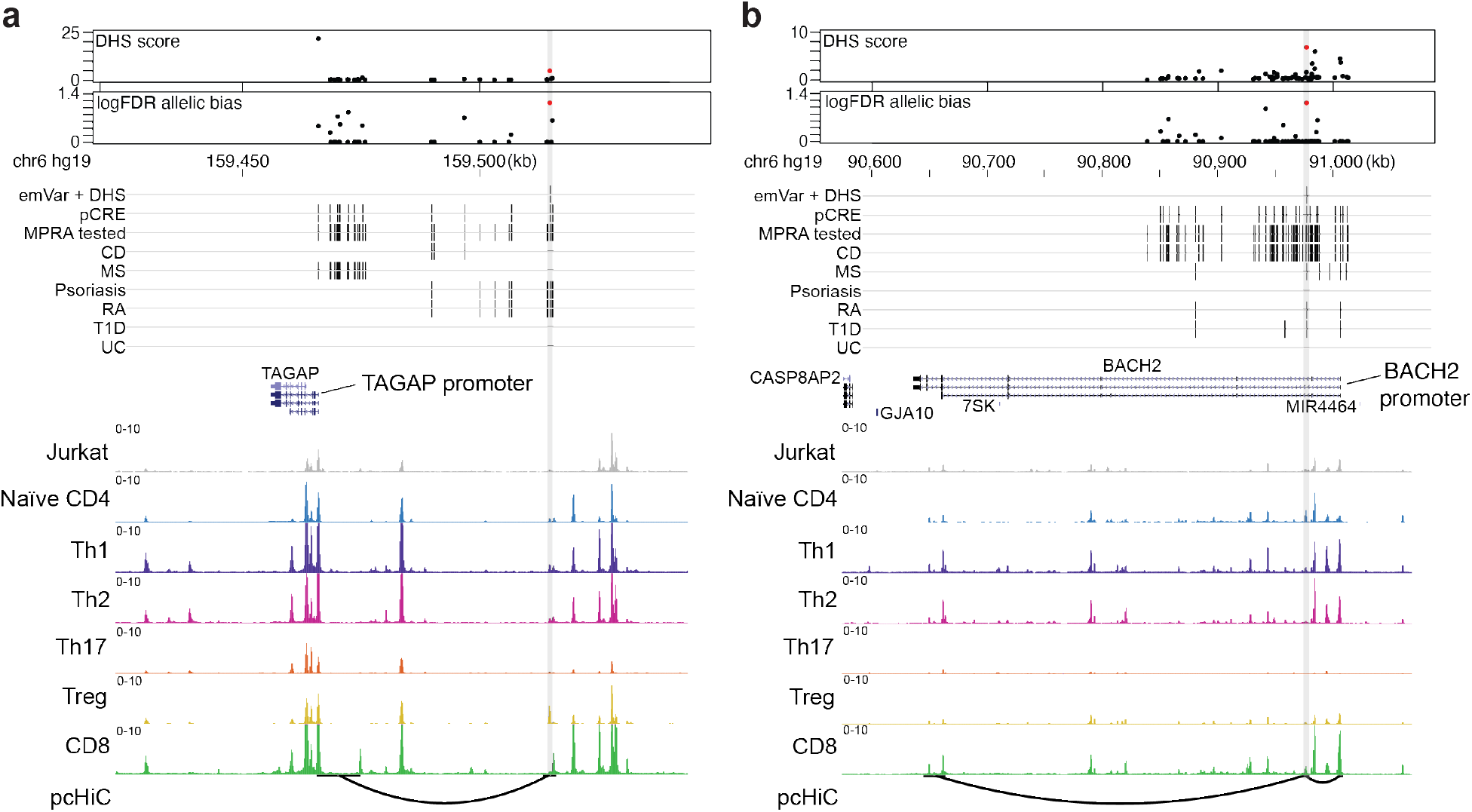
Putative causal variants in a *BACH2* intron and upstream of *TAGAP*. **a** and **b**) Dotplot (top) showing DHS signal (DHS score) and statistical significance of allelic bias (log_10_FDR of MPRA allelic bias) for MPRA variants in the region; all tested variants on haplotype (black), significant emVars in DHS (red dot). Position of variants that are emVars, pCREs, variants tested in MPRA, and disease-associated variants for CD, MS, psoriasis, RA, T1D, and UC from the GWAS Catalog^56^ (middle). Genes in the locus are shown along with chromatin accessibility profiles (from Jurkat and specific T cell subsets) and T cell promoter capture HiC (pcHiC^23^) loops anchored on the region containing the emVar. pcHiC loops in **(a)** are specific to naïve T cells; pcHiC loop in **(b)** is present in all T cell subsets and conditions tested. Gray line depicts position of the prioritized emVar with respect to all data types. Statistical significance of allelic biases in (**a**) and **(b)** were calculated using a paired Student’s two-sided *t*-test as described in Methods.

### An emVar in accessible chromatin reduces *BACH2* expression

We further characterized rs729282038, as it displayed one of the strongest allelic biases in reporter activity in the MPRA (Fig. 2c). We first validated our MPRA results for rs72928038 using a luciferase assay in Jurkat T cells. We cloned a construct that contained either allele of rs72928038 centered in 300 bp of native genomic sequence upstream of the *BACH2* promoter. We found that the risk allele (A) had lower luciferase activity (Supplementary Fig. 9a). There were two other statistically fine-mapped variants in the locus, rs10944479 (PICS probability 0.0458 in MS GWAS) and rs6908626 (posterior probability 0.0894 in MS GWAS) (in comparison, rs72928038 had posterior probability of 0.865 for MS GWAS). Neither of these variants were found to have allelic bias in the MPRA. These data suggest that, rs72928038 is the only variant in the credible set that alters regulatory activity (Fig. 4a).

**Fig. 4.**
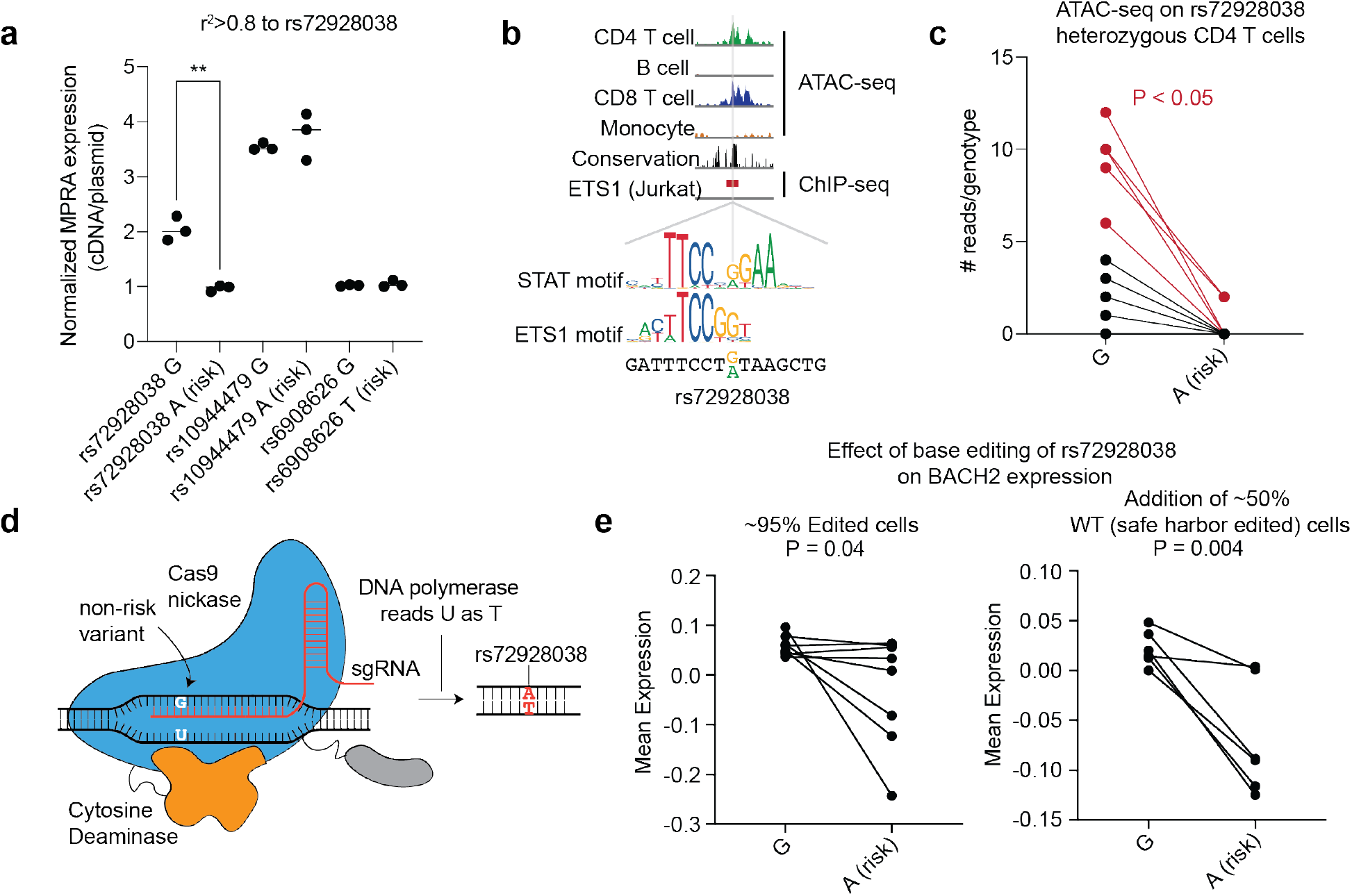
Base editing of the *BACH2* emVar (rs72928038) reduces *BACH2* expression. **a**) MPRA reporter expression of credible set variant alleles at the *BACH2* locus (n=3 independent replicates). **b**) ATAC-seq profiles in CD4 T cells, B cells, CD8 T cells, and monocytes, vertebrate conservation, and ETS1 ChIP-seq^57^ at the site of rs72928038 (top). STAT and ETS1 TF motifs at the site of rs72928038 (bottom). **c**) ATAC-seq reads overlapping rs72928038 in CD4 T cells from heterozygous healthy individuals (10 genotyped individuals);. 5 of the 10 individuals (marked red) had a significant difference (at *P* < 0.05) in number of reads between reference and alternate alleles according to a binomial test. **d**) Schematic of base editing rs72928038 using the evoCDAmax cytosine base editor. **e**) PrimeFlow mean expression of *BACH2* in cells containing the rs72928038 non-risk (G) and base-edited risk (A) allele with rs72928038 base-edited cells alone (left; 8 independent replicates) and when combined with cells that were edited at a safe harbor locus (right; 6 independent replicates). For **(a),** central tendency is shown as median and all points are plotted to show dispersion. For **(e),** central tendency is shown as mean and all points are plotted to show dispersion. P values determined by Student’s two-sided t-test (**a**); Binomial test (**c);** Student’s one-sided t-test (**e**).

rs729282038 is an eQTL specifically in naïve CD4, naïve CD8, and naïve regulatory T cells (but not other immune or T cell types) with the risk allele (A) associated with lower expression of *BACH2*^30^. If rs72928038 acts through modulating enhancer activity in T cells to alter *BACH2* expression, one may expect the variant to reside in accessible chromatin of T cells, but not of other cell types. Indeed, we found rs72928038 is located in accessible chromatin specifically in T cells, and not in B cells or monocytes (Fig. 4b). To assess differences in chromatin accessibility between alleles, we surveyed CD4 T cells from healthy donors who were heterozygous at rs72928038 and observed the non-risk allele (G) to be preferentially present in accessible chromatin (Fig. 4c; Supplementary Table 11). The 328 bp region surrounding rs72928038 is annotated as a candidate cis-regulatory element by ENCODE (EH38E2485452)^31^ and the risk allele is predicted to disrupt binding motifs for ETS or STAT family TFs (Fig. 4b; Supplementary Table 5). Furthermore, based on published promoter-capture HiC data, the region surrounding rs72928038 physically interacts with the *BACH2* promoter specifically in naïve T cells, but not other immune cell types (Supplementary Fig. 9b), suggesting that the risk allele of rs72928038 regulates *BACH2* expression in T cells through reducing cis-regulatory activity.

We next sought more direct functional evidence for the role of rs72928038 in altering *BACH2* expression. To do this, we used a cytosine base editor along with a guide RNA targeting rs72928038 to introduce the risk allele into the native genomic context in Jurkat T cells, which are homozygous for the non-risk allele (Methods)^32^ (Fig. 4d and Supplementary Fig. 10). Within the pool of nucleofected cells, we found 95% edited cells, with a range of bases edited in the editing window including specific edits of the variant of interest. To assess the effect of the risk variant on *BACH2* expression, we used PrimeFlow, which uses *in situ* hybridization of antisense probes to the *BACH2* transcript, followed by signal amplification, fluorescent labeling, and fluorescence-activated cell sorting to isolate cells that have either high or low *BACH2* expression (Supplementary Fig. 10)^6,7^. For each bin, we sequenced the rs72928038 region and compared the prevalence of amplicons containing the edited risk allele to those without edits, finding that the risk variant reduces *BACH2* expression (Fig. 4e, left). However, since unedited cells were only 5% of the cell population, the estimated expression levels of *BACH2* in the WT cells were variable leading to unstable estimates of effect. To address this, we created a second condition with cells nucleofected with base editor and either a guide RNA targeting rs72928038 or a safe harbor sequence, and then combined these cells at a 50/50 ratio, again finding that the base-edited risk variant confers reductions in *BACH2* expression (Fig. 4e, right; Supplementary Fig. 10). Together, these experiments show that the rs72928038 risk allele reduces the expression of *BACH2* in a human T cell line.

### Deletion of the orthologous non-coding element containing rs72928038 in mice leads to reduced expression of T cell stemness genes

To investigate the phenotypic effects of the regulatory region containing rs72928038 in primary naïve T cells, ideally one would isolate naïve T cells from secondary lymphoid organs, where they reside in T cell zones awaiting activation by an antigen presenting cell. Since these secondary lymphoid organs are easily harvested from mice, we explored the use of mice to study the effect of rs72928038 on naïve T cells. We first assessed conservation between human and mouse at the site of the variant. Through synteny analysis of the locus between human and mouse, we found that the variant exists on mouse chromosome 4 within an intron of *Bach2*, similar to its position with respect to *BACH2* in the human genome (Fig. 5a). The 328 bp human cCRE containing rs72928038 is 51.2% conserved between species, with especially high conservation in the 16 bps surrounding rs72928038 (Fig. 5a). Additionally, at the orthologous region, mouse T cells have accessible chromatin, H3K27ac deposition, and we found both ETS1 and STAT TFs bind (Supplementary Fig. 11a)^33^, consistent with the epigenetic profile observed in human T cells. Based on these findings, we created a mouse line containing an 18bp deletion of the non-coding region overlapping the variant using CRISPR-mediated genome editing (Bach2^18del^; Fig. 5b, Supplementary Fig. 11a).

**Fig. 5.**
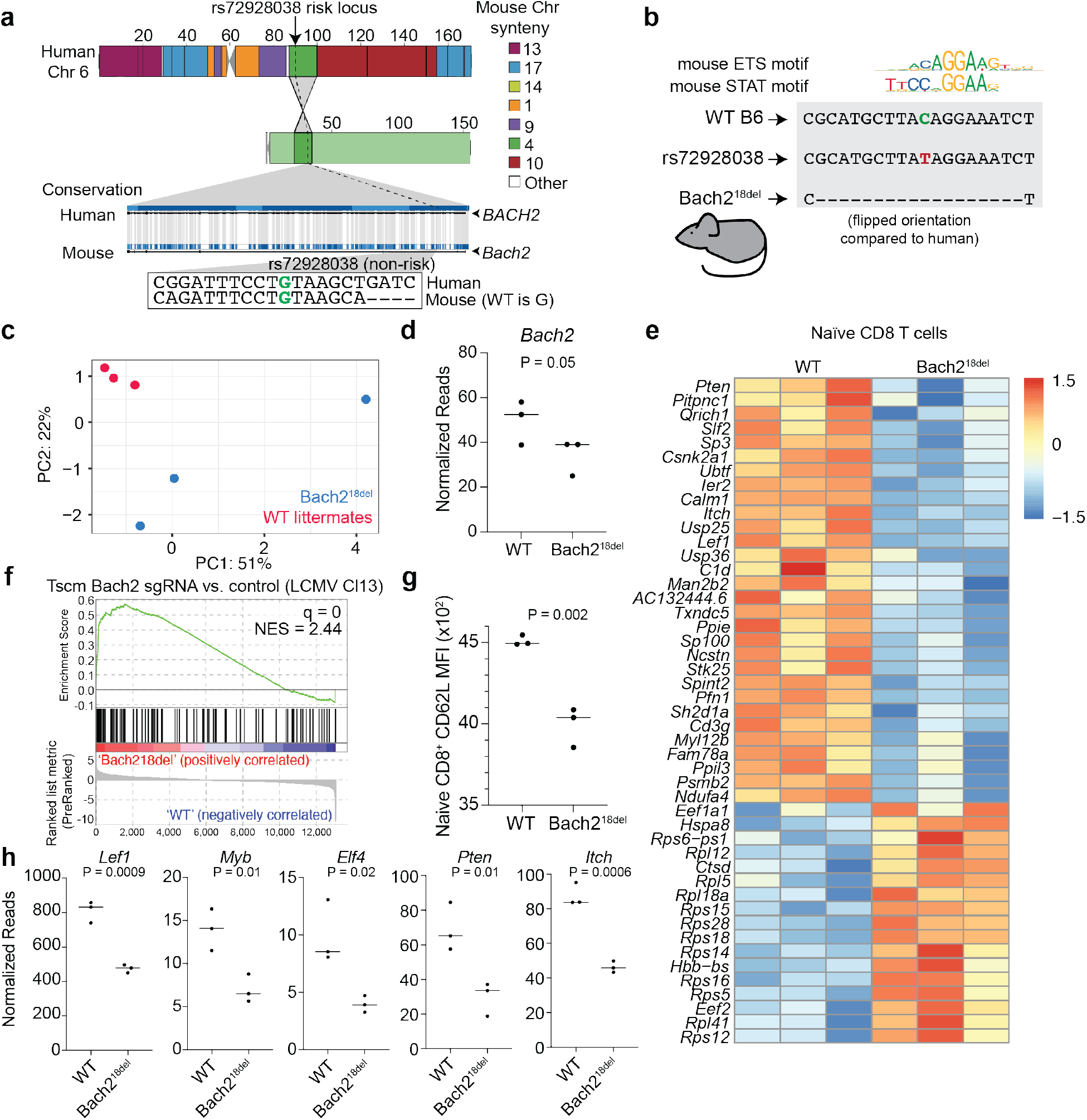
Naïve T cells from mice with a deletion overlapping orthologous rs72928038 have reduced transcriptional features of stemness. **a**) Synteny analysis of rs72928038 between human and mouse. Location of rs72928038 (arrow and dotted line) on human chromosome 6 and mouse chromosome 4 (top) with colors indicating mouse chromosome synteny (see key). Conservation of human *BACH2* with mouse *Bach2*, with the location of rs729280378 noted (dotted line), with inset showing conserved sequence between human and mouse at the site of rs72928038 (bottom). **b**) Schematic of Bach2^18del^ mutation. **c**) Principal components 1 and 2 from gene expression analysis of naïve CD8 T cells from Bach2^18del^ and their WT littermates. **d**) *Bach2* gene expression within naïve CD8 T cells from WT and Bach2^18del^ mice. **e**) Expression heatmap of differentially expressed genes (adjusted P value ≤ 0.05; calculated as described in Methods) between WT and Bach2^18del^ naïve CD8 T cells (n = 3 per genotype). **f**) GSEA showing enrichment of Bach2^18del^ vs. WT naïve CD8 T cells for a gene set derived from genes differentially expressed in *Bach2* guide RNA-targeted CD8 Tscm cells vs. empty vector Tscm cells. Full GSEA results are shown in Table S13. **g**) Mean fluorescence intensity of CD62L surface expression on naïve WT and Bach2^18del^ CD8 T cells. **h**) Sample genes differentially expressed between WT and Bach2^18del^ mice in naïve CD8 T cells (c-h, n = 3 per genotype). P values determined by Student’s one-sided t-test (**d**); Student’s two-sided t-test (**g** and **h**). For (**d**), (**g**), and (**h**), central tendency shown as median and all points are plotted to show dispersion. Normalized enrichment score (NES) in **(f)** was calculated based on observed enrichment as compared to enrichments from permuted data as previously described and statistical significance shown as the false discovery rate (q)^58^.

Using these mice, we performed experiments to determine if primary mouse naïve T cells containing the deletion had reduced *Bach2* expression and altered expression of other important genes that play a role in T cell biology. *Bach2*-ablated mice have previously been shown to have aberrant CD8 T cell activation^8^. To assess whether deletion of the variant alters naïve CD8 T cell phenotypes in mice, we sorted naïve CD8 T cells from Bach2^18del^ and WT mice and analyzed their transcriptomes via BRB-seq^34^. We found Bach2^18del^ naïve CD8 T cells to have altered transcriptional programs according to principal components analysis (PCA) and reduced *Bach2* expression compared to WT littermates (Fig. 5c and d). Through differential expression analysis, we found 47 differentially expressed genes (Fig. 5e; Supplementary Table 12). Genes more highly expressed in Bach2^18del^ naïve CD8 T cells were enriched for KLRG1^lo^ effector CD8 T cell gene sets (Supplementary Table 13), as well as for a gene set in which CD8 T stem-like memory cells (Tscms) were perturbed with CRISPR/Cas9 targeted against *Bach2* (Fig. 5f; Supplementary Fig. 11b)^10^. We found 66% of the differentially expressed genes from Bach2^18del^ naïve CD8 T cells have the same directionality in *Bach2* guide RNA targeted Tscms (Supplementary Fig. 11c). Similar to *Bach2*-perturbed Tscms, Bach2^18del^ naïve CD8 T cells had significantly reduced CD62L surface expression (Fig. 5g), concomitant with a reduction in *Lef1* and *Myb* expression (Fig. 5h); these TFs are required to maintain stemness of naïve T cells and Tscms, and are downregulated during effector T cell differentiation^10,35–37^. In addition, Bach2^18del^ naïve CD8 T cells showed a reduction in *Elf4* and a significant upregulation of ribosomal protein mRNAs, both indications of early T cell stimulation^38,39^ (Fig. 5e and h). Bach2^18del^ naïve CD8 T cells also had reduced expression of *Pten* and *Itch*, both negative regulators of signaling that are required for suppressing effector T cell differentiation^40,41^ (Fig. 5h). Thus, deletion of the orthologous non-coding region containing rs72928038 in mice leads to reduced features of naïve T cell stemness and indications of early T cell activation.

## DISCUSSION

Identifying mechanisms that drive genetic risk for autoimmunity and other complex phenotypes remains a substantial challenge. Most autoimmune genetic associations have many variants in tight LD to the lead variant^3^, and the majority of these variants have not been functionally characterized using a systematic approach within disease-relevant cell types. Here, using a combination of MPRA and T cell chromatin accessibility, we identify 60 variants associated with five autoimmune diseases that enrich 57.8-fold for causal variants according to statistical fine-mapping. Collectively, these data demonstrate that this combination of methods serves as a robust prioritization scheme for identifying causal variants for disease associations. Many GWAS loci with T-GWAS emVars are near genes that regulate T cell function, or those that are known to dysregulate T cells in the context of autoimmune disease. One of the variants with high allelic bias in MPRA was rs72928038, a variant in an intron of *BACH2*. We found the rs72928038 risk allele was less present in chromatin accessibility data from T cells harvested from humans heterozygous for rs72928038, and base editing of the rs72928038 risk allele in a human T cell line reduced *BACH2* expression. We also engineered mice that have an 18 bp deletion overlapping orthologous rs72928038 (Bach2^18del^), and found that their naïve CD8 T cells had reduced *Bach2* expression, as well as reduced expression of transcriptional regulators of T cell stemness and indications of early T cell activation.

By applying MPRA, we substantially enriched for statistically fine-mapped GWAS variants, an enrichment that was further magnified when combining these data with chromatin accessibility data from T cells. Combining MPRA and accessible chromatin data was an effective strategy, possibly because chromatin accessibility, which provides an endogenous measure of cis-regulatory activity from relevant cell types, acts as a stringency filter for MPRA, which is plasmid-based. This strategy was further supported by an increase in sensitivity for identifying credible set variants from 18.4-19.7% to 23.1-25.5% while maintaining a specificity of 81.1-91.7%. Other prioritization methodologies could also be applied in tandem such as allele-specific ATAC-seq^12^, CRISPR-inhibition^7,42^, and SELEX^43^, among others. However, requiring a variant to score for multiple methodologies may substantially increase type II error, as different methods tend to test different genomic features and have variable signal-to-noise ratios^6^. Thus, combining data from orthogonal tests of variant action with high signal-to-noise ratios, such as MPRA and accessible chromatin, could provide a reasonable balance between sensitivity and specificity^6^. Because perturbational and statistical fine-mapping are still imperfect approaches, discovery of causal variants still requires further mechanistic evaluation, ideally within systems that recapitulate the (patho)-physiological environment of the disease.

While we discovered emVars for ∼31% of GWAS loci studied, there are a variety of reasons why many loci did not contain an identified emVar. We found emVars to be enriched in TSS regions, thus this methodology may have increased sensitivity for variants that alter promoter activity. We performed the MPRA in unstimulated conditions, although variants may disrupt TFs that are downstream of signaling cascades following T cell stimulation or differentiation into specific effector cell subsets (e.g., Th1, Th2, Th17, Treg, Tfh). Stimulation with various ligands in eQTL studies has been crucial for identifying variants that were otherwise inactive at baseline^44–46^. Other cell types also likely play a role in these autoimmune diseases; beta cell accessible chromatin is enriched for T1D-associated variants, especially after stimulation with pro-inflammatory cytokines^47^, B cell accessible chromatin enriches for MS-associated variants^48^, skin cell accessible chromatin enriches for psoriasis^49^, and intestinal CAGE data enriches for IBD-associated variants^50^. Variants in GWAS loci may have roles beyond disrupting cis-regulatory elements, such as coding mutations, altering the activity of untranslated regions (UTRs), or promoting alternative splicing. These actions are unlikely to be identified by MPRAs designed to test how variants modulate regulatory region activity, but alternative massively parallel methodologies have been created to address how variants may alter UTR function and alternative splicing^51,52^. Thus, applying the prioritization scheme of MPRA + accessible chromatin and other methodologies to a wider range of cell types and stimulation conditions could unveil additional likely causal variants.

Using our MPRA and accessible chromatin prioritization scheme, we found variants in GWAS loci that were highly relevant to T cell biology, including rs72928038 in the *BACH2* locus. We selected rs72928038 for further mechanistic studies due to its high allelic bias in MPRA and the strength of genetic and epigenetic evidence supporting the variant. Through base editing the variant in Jurkat T cells and deleting the orthologous region surrounding the variant in mice, we found rs72928038 and the regulatory region in which it is contained altered T cell *BACH2* expression. Similar to *Bach2-*deficient CD8 stem-like memory cells from a separate study^10^, we observed that Bach2^18del^ naïve CD8 T cells have reductions in stem-associated TFs *Lef1*, *Myb*, antiproliferative TF *Elf4*, signaling attenuators *Pten* and *Itch*, and a reduction in surface expression of CD62L. We also found an increase in expression of ribosomal genes, indicative of early T cell stimulation^39^. Consistent with increased activation, we saw enrichments of effector CD8 T cell gene programs. Collectively these data suggest that the Bach2^18del^ naïve CD8 T cells may have a reduced threshold for activation. However, Bach2^18del^ naïve CD8 T cells do not appear to have phenotypes of fully differentiated effector cells (e.g., increased expression of *Gzmb*, *Klrg1* and *Cd44*), possibly due to only partial *Bach2* reduction mediated by removing only a single regulatory element as opposed to deletion of the gene. Indeed several TFs have been noted to act in a graded manner to promote transcription and cell fate during the differentiation of CD8 T cells^53,54^, and mice heterozygous for the deletion of *Bach2* show intermediate effects on T cell differentiation between WT and homozygous mice^9^. Other than naïve T cells, further dissection will be required to understand the effects of rs72928038 on subsets of T cells that are known to require BACH2 for fate determination, such as Tscms, memory T cells, and Tregs^8–10^. Thus, organismic models, such as the Bach2^18del^ mice, provide rare insight into the physiological effects of variants and their regulatory elements within living systems.

In summary, this work provides a scalable and high-yield prioritization scheme to identify likely causal variants at high specificity. We find 60 likely causal variants that have significant evidence for acting in T cells, and direct evidence of a variant that reduces *Bach2* expression and transcriptional hallmarks of naïve T cells and T cell stemness. Together, this work demonstrates a clear path for addressing the long-term obstacle of defining causal variants for complex traits and their effects on gene regulation and cellular and organismal functions.

## METHODS

### Cell lines

For MPRA, luciferase, and base editing experiments, we used low passage aliquots of the Jurkat T cell line (ATCC TIB-152™), maintaining the culture under 20 passages. Cells were grown at 37 °C maintaining cultures between 1×10^5^ and 1×10^6^ cells per mL.

### Study subjects

The study was performed in accordance with protocols approved by the institutional review board at Partners (Brigham and Women’s Hospital, Massachusetts General Hospital, Dana-Farber Cancer Institute, Boston, USA) and Broad Institute (USA) Research Ethics Committee, as well as the Feinstein Institute for Medical Research, Northwell Health institutional review board (Manhasset New York, USA). All donors provided written informed consent for the genetic research studies and molecular testing. Healthy donors were recruited from the Sisters of Lupus Erythematosus patients (SisSLE) Research Study based in Manhasset, NY, and the Boston-based PhenoGenetic project, a resource of healthy subjects.

### GWAS data

Lead SNPs were obtained from GWAS for T1D^20^, RA^22^, psoriasis^21^, IBD^17^, and MS^18,19^. We collected 578 GWAS lead SNPs from these studies, representing 531 distinct GWAS loci. We identified all proxy SNPs (r^2^ ≥ 0.8) for each lead SNP based on 1000 Genomes Phase 3 European subset. Proxy SNPs were identified using PLINK v1.90b3.32^60^ (www.cog-genomics.org/plink/2.0/) with parameters --r2 --ld-window-kb 2000 --ld-window 999999 --ld-window-r2 0.8. There were 20792 total proxy SNPs across the 578 GWAS loci (18324 unique proxy SNPs across these 531 distinct GWAS loci).

### MPRA

MPRA oligo synthesis and cloning was adapted from refs ^6,13^. Each allele was tagged with an average of ∼1,000 DNA barcodes. Oligos were synthesized by Agilent Technologies containing 170 bp of genomic context and 15 bp of adapter sequence at either end (5′-ACTGGCCGCTTGACG[170 bp oligo]CACTGCGGCTCCTGC-3′; Supplementary Table 14; 200 bp total). 20 bp barcodes and additional adapter sequences were added by performing 28 emulsion PCR reactions, each 50 μL in volume containing 1.86 ng of oligo, 25 μL of Q5 NEBNext MasterMix (NEB, M0541S), 1 unit Q5 HotStart polymerase (NEB, M0493S), 0.5 μM MPRA_v3_F and MPRA_v3_20I_R primers (Supplementary Table 14) and 2 ng BSA (NEB, B9000). PCR master mix was emulsified by vortexing with 220 μL Tegosoft DEC (Evonik), 60 μL ABIL WE (Evonik) and 20 μL mineral oil (Sigma, M5904) per 50 μL PCR reaction at 4 °C for 5 min. 50 μL of emulsion mixture was added to each well of a 96-well plate and cycled with the following conditions; 95 °C for 30 s, 15 cycles of (95 °C for 20 s, 60 °C for 10 s, 72 °C for 15 s), 72 °C for 5 min. Amplified emulsion mixture was broken and purified by adding 1 mL of 2-butanol (VWR, AA43315-AK), 50 μL of AMPure XP SPRI (Beckman Coulter, A63881) and 80 μL of binding buffer (2.5 M NaCl, 20% PEG-8000) per 350 μL of emulsion mix and vigorously vortexed, followed by incubation for 10 min at room temperature. Broken emulsion/butanol mixture was spun at 2900 × *g* for 5 min and the butanol phase was discarded. The aqueous phase was placed on a magnetic rack for 20 min prior to aspiration. Remaining beads were washed once with 2-butanol, three times with 80% EtOH and eluted in EB (Qiagen, 19086) to yield the barcoded oligo pool.

To create our mpraΔorf library, barcoded oligos were inserted into SfiI digested pGL4.23ΔxbaΔluc by Gibson Assembly (NEB, E2611) using 1.1 μg of oligos and 1 μg of digested vector in a 40 μL reaction incubated for 60 min at 50 °C followed by AMPure XP SPRI purification and elution in 20 μL of EB. Half of the ligated vector was then transformed into 200 μL of Endura^TM^ ElectroCompetent *E. coli* (Lucigen, 60242-2) by electroporation (1.8 kV, 600 Ω, 10 μF). Electroporated bacteria were immediately split into eight 1 mL aliquots of SOC (NEB, B9020S) and recovered for 1 h at 37 °C then independently expanded in 20 mL of LB supplemented with 100 μg/mL of carbenicillin (EMD, 69101-3) on a floor shaker at 37 °C for 6.5 h. After outgrowth, aliquots were pooled prior to plasmid purification (Qiagen, 12963). For each of the aliquots, we plated serial dilutions after SOC recovery and estimated a library size of ∼4 × 10^7^ CFUs, representing ∼1000 barcodes per allele.

To insert minP and GFP ORF, 20 μg of mpra:Δorf plasmid was linearized with AsiSI (NEB, R0630S) and 1x CutSmart buffer (NEB, B7204S) in a 500 μL volume for 3.5 h at 37 °C, followed by bead inactivation for 20 m at 80 °C and SPRI cleaning. An amplicon containing minP, the GFP open-reading frame, and a partial 3′ UTR was then inserted by Gibson assembly using 10 μg of AsiSI linearized mpraΔorf plasmid, 33 μg of the minP/GFP amplicon in 400 μL of total volume for 90 min at 50 °C followed by a 1.5× beads/sample SPRI purification. The total recovered volume was digested a second time to remove remaining uncut vectors by incubation with AsiSI in a 100 μL reaction for 6 h at 37 °C followed by Ampure XP purification and elution with 55 μL of Buffer EB.

10 μL of the mpra:minP:gfp plasmid was electroporated (1.8 kV, 600 Ω, 10 μF) into 200 μL of Endura cells. Electroporated bacteria was split across six tubes and each recovered in 2 mL of SOC for 1 h at 37 °C then added to 500 mL of LB with 100 μg/mL of carbenicillin and grown for 9 h at 37 °C prior to plasmid purification (Qiagen, 12991). The plasmid prep was then normalized to 1 μg/μL to generate the final mpra:minP:gfp library used for transfection delivery.

For all transfections, cells were grown to a density of ∼1 × 10^6^ cells/mL, and 1 × 10^8^ cells were used for each experiment. Cells were collected by centrifugation at 300 × *g* and eluted in 1 mL of RPMI with 100 μg of mpra:minP:gfp library. Electroporation was performed in 100 μL volumes with the Neon transfection system (Life Technologies) applying three pulses of 1600V for 10 ms each into Jurkat T cells. Using separate control transfections, we achieved transfection efficiencies of 40–60% for all replicates. Cells were allowed to recover in 200 mL in RPMI with 15% FBS for 24 h before being collected by centrifugation, washed once with PBS, collected and frozen at −80 °C.

Total RNA was extracted from cells using QIAGEN Maxi RNeasy (QIAGEN, 75162) following the manufacturer’s protocol including the on-column DNase digestion. A second DNase treatment was performed on the purified RNA using 5 μL of Turbo DNase (Life Technologies, AM2238) with buffer, in 750 μL of total volume for 1 h at 37 °C. The digestion was stopped with the addition of 7.5 μL 10% SDS and 75 μL of 0.5 M EDTA followed by a 5-min incubation at 70 °C. The total reaction was then used for pulldown of GFP mRNA. Water was added to the DNase-digested RNA to bring the total volume to 898 μL to which 900 μL of 20X SSC (Life Technologies, 15557-044), 1800 μL of Formamide (Life Technologies, AM9342) and 2 μL of 100 μM biotin-labeled GFP probe (GFP_BiotinCapture_1-3, IDT, Supplementary Table 14) were added and incubated for 2.5 h at 65 °C. Biotin probes were captured using 400 μL of pre-washed Streptavidin beads (Life Technologies, 65001) eluted in 500 μL of 20X SSC. The hybridized RNA/probe bead mixture was agitated on a nutator at room temperature for 15 min. Beads were captured by magnet and washed once with 1× SSC and twice with 0.1× SSC. Elution of RNA was performed by the addition of 25 μL water and heating of the water/bead mixture for 2 min at 70 °C followed by immediate collection of eluent on a magnet. A second elution was performed by incubating the beads with an additional 25 μL of water at 80 °C. A final DNase treatment was performed in 50 μL total volume using 1 μL of Turbo DNase with addition of the buffer incubated for 60 min at 37 °C followed by inactivation with 1 μL of 10% SDS and purification using RNA clean SPRI beads (Beckman Coulter, A63987).

First-strand cDNA was synthesized from half of the DNase-treated GFP mRNA with SuperScript III and a primer specific to the 3′ UTR (MPRA_v3_Amp2Sc_R, Supplementary Table 14) using the manufacturer’s recommended protocol, modifying the total reaction volume to 40 μL and performing the elongation step at 47 °C for 80 min. Single-stranded cDNA was purified by SPRI and eluted in 30 μL EB.

To minimize amplification bias during the creation of cDNA tag sequencing libraries, samples were amplified by qPCR to estimate relative concentrations of GFP cDNA using 1 μL of sample in a 10 μL PCR reaction containing 5 μL Q5 NEBNext master mix, 1.7 μL SYBR Green I diluted 1:10,000 (Life Technologies, S-7567) and 0.5 μM of TruSeq_Universal_Adapter and MPRA_Illumina_GFP_F primers (Supplementary Table 14). Samples were amplified with the following qPCR conditions: 95 °C for 20 s, 40 cycles (95 °C for 20 s, 65 °C for 20 s, 72 °C for 30 s), 72 °C for 2 min. The number of cycles for sample amplification was 1−*n* (the number of cycles it took for each sample to pass the threshold) from the qPCR. To add Illumina sequencing adapters, 10 μL of cDNA samples and mpra:minP:gfp plasmid control (diluted to the qPCR cycle range of the samples) were amplified using the reaction conditions from the qPCR scaled to 50 μL, excluding SYBR Green I. Amplified cDNA was SPRI purified and eluted in 40 μL of EB. Individual sequencing barcodes were added to each sample by amplifying the entire 40 μL elution in a 100 μL Q5 NEBNext reaction with 0.5 μM of TruSeq_Universal_Adapter primer and a reverse primer containing a unique 8 bp index (Illumina_Multiplex, Supplementary Table 14) for sample demultiplexing post-sequencing. Samples were amplified at 95 °C for 20 s, six cycles (95 °C for 20 s, 64 °C for 30 s, 72 °C for 30 s), 72 °C for 2 min. Indexed libraries were SPRI purified and pooled according to molar estimates from Agilent TapeStation quantifications. Samples were sequenced using 1 × 30 bp chemistry on a NextSeq 2000 (Illumina).

To determine oligo/barcode combinations within the MPRA pool, Illumina libraries were prepared from the mpraΔorf plasmid library by performing five separate amplifications with 200 ng of plasmid in a 100 μL Q5 NEBNext PCR reaction containing 0.5 μM of TruSeq_Universal_Adapter and MPRA_v3_TruSeq_Amp2Sa_F primers (Supplementary Table 14) with the following conditions: 95 °C for 20 s, 6 cycles (95 °C for 20 s, 62 °C for 15 s, 72 °C for 30 s), 72 °C for 2 min. Amplified material was SPRI purified using a 0.6 × bead/sample ratio and eluted with 30 μL of EB. Sequencing indexes were then attached using 20 μL of the eluted product and the same reaction conditions as for the tag-seq protocol, except the number of enrichment cycles was lowered to 5. Samples were molar pooled and sequenced using 2 × 150 bp chemistry on an Illumina NextSeq 2000.

### MPRA analysis

Barcode sequencing results from the MPRA were analyzed as previously described^13^. Briefly, the sum of the barcode counts for each oligo was provided as input to DESeq2 and replicates were median normalized followed by an additional normalization of the RNA samples to center the RNA/DNA activity distribution over a log_2_ fold change of zero^72^. Oligos showing differential expression relative to the plasmid input were identified by modeling a negative binomial distribution with DESeq2 and applying a false discovery rate (FDR) threshold of 1%. For sequences that displayed significant MPRA activity, a paired two-sided Student’s *t*-test was applied on the log-transformed RNA/plasmid ratios for each experimental replicate to test whether the reference and alternate allele had similar activity. An FDR threshold of 10% was used to identify SNPs with a significant difference in MPRA activity between alleles (emVars).

### Genomic data integration and enrichment analyses

DHS data across 733 samples was obtained from ^4^. Pre-processed DHS peaks lifted to hg19 were downloaded from https://zenodo.org/record/3838751#.X_IA7-lKg6U. For each of 261 unique cell types, DHS peaks were merged across replicates of the same cell type. To assess enrichment between DHS sites for each cell type and MPRA data, we counted the number of MPRA variants with regulatory activity (i.e., pCRE) overlapping the genomic interval bed file for each cell type. We then constructed a 2 x 2 contingency table based on whether the MPRA SNP showed regulatory activity (i.e., pCRE) or had no regulatory activity, and whether the SNP intersected a genomic interval or not. P values were calculated based on a two-sided Fisher’s exact test. Multiple testing correction was performed using Bonferroni correction by taking an alpha of 0.05 divided by the number of unique cell types tested (n=261).

Histone ChIP-seq data were obtained from the ENCODE project for all available human CD4 positive alpha-beta T cell samples. For each cell type, all available ChIP-seq peak call sets for H3K4me1, H3K4me3, H3K27ac, and H3K36me3 aligned to hg19 were downloaded in bed file format. If multiple replicates were available, peaks call sets were merged using the merge function in BEDTools v2.26.0^61^. Human CAGE-based enhancer sequences were downloaded from https://fantom.gsc.riken.jp/5/datafiles/latest/extra/Enhancers/ in bed file format. chromHMM annotations in the 18 chromatin state model were obtained for primary T cells from peripheral blood (E034) from the Roadmap Epigenomics Project (https://egg2.wustl.edu/roadmap/data/byFileType/chromhmmSegmentations/ChmmModels/core_K27ac/jointModel/final/) in bed file format. To assess enrichment between each of these genomic datasets and MPRA data, we counted the number of emVar SNPs overlapping the genomic interval bed file. We then constructed a 2 x 2 contingency table based on whether the SNP showed MPRA activity (i.e., emVar) or had no MPRA activity, and whether the SNP intersected a genomic interval or not. P values were calculated based on a two-sided Fisher’s exact test. Multiple testing correction was performed using Bonferroni correction by taking an alpha of 0.05 divided by the number of genomic annotations tested.

### Transcription factor enrichment

To calculate the predicted effect of each MPRA variant on TF binding, we applied motifbreakR version 2.4.0^62^. For each single nucleotide substitution in the MPRA data, we calculated predicted TF binding scores for the reference and alternate alleles. We used the sum of log probabilities approach in motifbreakR, applied to all TF position-weighted matrices in HOCOMOCO v10^63^. All MPRA SNPs with a difference in TF binding scores between the reference and alternate alleles at *P* < 1×10^−5^ were considered to be significant.

To assess overlaps between MPRA SNPs and TF binding motifs derived from ChIP-seq, we applied HOMER v4.10^64^. For each emVar, we generated sequences in a ± 100 bp window around the emVar SNP. We then generated background sequences in a ± 100 bp window around all variants tested in the MPRA. To calculate enrichments, we applied findMotifGenome.pl in HOMER with -size 200; all other default parameters were used. Enrichments were tested against the library of all known TF motifs available in HOMER.

### ATAC-seq skew and QTL enrichment

To assess enrichment of MPRA SNPs at sites of allelic skew in open chromatin regions, we downloaded significant allelic skew SNPs from Calderon et al.^12^. We used all available unstimulated hematopoietic cell types. For SNPs with evidence of skew across multiple samples or cell types, we summed the reference and alternate allele counts for that SNP across the cell types. To assess enrichment of MPRA SNPs at ATAC-QTLs, we downloaded primary T-cell ATAC-QTLs from^65^.

### *In silico* predictions of effect of regulatory activity

We applied deltaSVM v1.3^66^ to predict the effect of MPRA SNPs on regulatory activity. This method uses a classifier (gkm-SVM) to encode cell-specific regulatory sequence vocabularies, and then subsequently deltaSVM quantifies the effect of a SNP as the change in gkm-SVM score). To apply deltaSVM, we downloaded pre-computed gkm-SVM weights derived from ENCODE2 enhancers in naïve CD4 T cells. We then used deltasvm.pl from the software developers to calculate deltaSVM scores for each emVar. Default parameters were used throughout.

### PICS fine-mapping and enrichment analyses

For each GWAS locus, PICS^1^ was applied to all SNPs in LD (r^2^ ≥ 0.8) to the lead SNP at based on the 1000 Genomes Phase 3 European subset using PLINK v1.90b3.32 with parameters --r2 -- ld-window-kb 2000 --ld-window 999999 --ld-window-r2 0.8. GWAS association P values for lead SNPs were obtained from the EMBL GWAS catalog^56^ on August 10, 2020^56^. If the same lead SNP was seen multiple times in the GWAS catalog for either the same disease or multiple diseases, the most significant lead SNP P value was used. Given long-range patterns of high LD, we excluded the human MHC locus (chr6:29691116-33054976 in hg19) and excluded any lead SNP where the most significant GWAS association P value did not reach 5 x 10^−8^. In total, 512 GWAS loci were analysed. PICS fine-mapping posterior probabilities were calculated using a custom PERL script. Of note, in the scenario where a lead SNP was seen multiple times (either across the same disease or shared by different diseases), all proxy SNPs to the lead SNP were assigned based on the most significant lead SNP association P value, and PICS probabilities were calculated for both this lead SNP and its proxies.

We defined SNPs as being statistically fine-mapped based on either if it had a posterior probability greater than a given threshold, or if it was in a fine-mapping credible set. We used posterior probability thresholds of ≥ 0.01, 0.05, 0.1, 0.2, 0.3, or 0.5. We also calculated credible sets of fine-mapping variants. An X% credible set is expected to contain the true causal variant X% of the time. To generate credible sets, we greedily summed up the highest fine-mapping posterior probabilities at each locus until reaching a cumulative X%. We further required all credible set variants to have a fine-mapping posterior probability ≥ 0.01.

To calculate enrichment of MPRA emVars in PICS fine-mapped SNPs, we determined for each MPRA variant whether it was a PICS statistically fine-mapped SNP (i.e., had a PICS probability greater than a given threshold) and whether the fine-mapped SNP showed MPRA activity (i.e., emVar). From this, we constructed 2 x 2 contingency tables. We then calculated variant-level enrichment (*E_v_*), represented as a risk ratio:

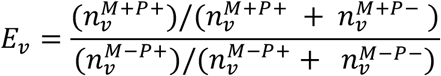

Here, *n_v_* refers to the number of variants. Superscript *M* refers to whether the variant is an emVar (*M* + if emVar, *M* − otherwise). Superscript *P*refers to whether the SNP is a statistically fine-mapped SNP by PICS (*P* + if PICS fine-mapped, *P* − otherwise). We can then construct a 2 x 2 contingency table with values 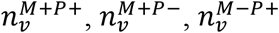, and 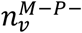. Statistical significance was calculated using a two-sided Fisher’s exact test based off this 2 x 2 contingency table.

We similarly calculated enrichment of MPRA variants overlapping T cell DHS sites in PICS statistically fine-mapped SNPs. We determined for each MPRA variant whether it directly overlapped a T cell DHS site and whether it was a PICS fine-mapped SNP (i.e., had a PICS probability greater than a given threshold). We then calculated variant-level enrichment (*E_v_*) represented as a risk ratio:

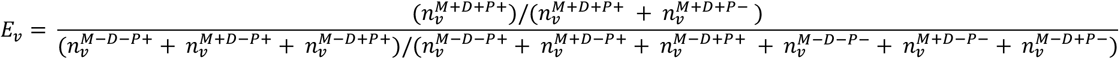

Here, superscript *D* refers to whether the variant is within a DHS site (*D* + if emVar, *D* − otherwise) and superscript *M* refers to whether the variant is an emVar (*M* + if emVar, *M* − otherwise). We can then construct a 2 x 2 contingency table with values 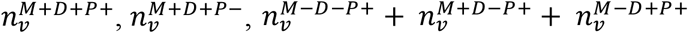, and 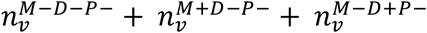. P values were calculated using a two-sided Fisher’s exact test.

We also calculated sensitivity and specificity of the MPRA using PICS fine-mapping as the benchmark “ground truth.” To do this, we tabulated for each locus, if there was a fine-mapped SNP (i.e., in X% credible set) and whether the fine-mapped SNP showed MPRA activity (i.e., emVar). We can then calculate sensitivity at the locus level (*SE_L_*):

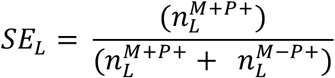

And specificity (*SP_L_*):

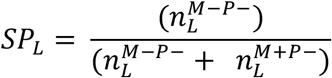

Here, *n_L_* refers to the number of GWAS loci. Superscript *M* refers to whether the variant is an emVar (*M* + if emVar, *M* − otherwise). Superscript *P*refers to whether the SNP is a statistically fine-mapped SNP by PICS (*P* + if PICS fine-mapped, *P* − otherwise).

### T1D GWAS fine-mapping enrichment analysis

For T1D GWAS, we obtained statistical fine-mapping credible sets from Onengut-Gumuscu et al.^20^. The main fine-mapping results from this study were used (Supplementary Table 1 of Onengut-Gumuscu et al.). We considered all T1D GWAS loci where an emVar was identified (n=11). For each MPRA variant in these loci, we then determined whether it was in the fine-mapping study credible set and had a fine-mapping posterior probability above a specified threshold. If a SNP was not part of the fine-mapping because it was present in 1000 Genomes Project but not ImmunoChip (SNPs designated by “proxy” from the T1D study authors), the SNP was assigned the fine-mapping posterior probability of its closest ImmunoChip proxy. To assess enrichments for MPRA in the T1D statistical fine-mapping data, we constructed a 2 x 2 contingency table based on whether the SNP did or did not show MPRA activity (i.e., emVar), and whether the SNP was a statistically fine-mapped variant. Enrichments and P values were calculated as described above.

### Visualization of GWAS loci

For visualization of gene tracks, bigWig files (Fig. 3a, 3b, Supplementary Fig. 8a and 8b) were downloaded from ENCODE. For cell types with multiple bigWig tracks, these were merged using bigWigMerge in the UCSC genome browser software suite^76^. bigWig tracks were then loaded into the UCSC genome browser (hg19). Track heights were adjusted to the maximum height of all tracks in a given viewing window. Gene transcripts are based on default UCSC genome browser gene annotations. To calculate the DHS score, the DHS sequencing depth in a ±10 bp window around each MPRA SNP was calculated using the multiBigwigSummary command in deepTools v3.5.0^77^ with default options. The DHS score plot shows the maximum DHS signal observed across each of the T cell types. PCHiC loops for Figs. 3a, 3b and Supplementary Fig. 9 were downloaded from https://www.chicp.org/. The visualized loops were selected from those with a CHiCAGO score ≥ 5 are shown.

### Luciferase Assay

Firefly luciferase reporter constructs (pGL4.24) were generated by cloning the 300 nucleotide genomic region centered on rs72928038 (rs72928038_luc_G and rs72928038_luc_A, Supplementary Table 14) of interest upstream of the *BACH2* promoter (Bach2_promoter_luc Supplementary Table 14) by using BglII and XhoI sites. The firefly luciferase constructs (500 ng) were nucleofected with a pRL-SV40 *Renilla* luciferase construct (50 ng) into 2 x 10^6^ Jurkat cells by using the Neon nucleofection system (Invitrogen) using the program 1600V, 3 pulses, 10ms. After 48 h, luciferase activity was measured by Dual-Glo Luciferase assay system (Promega) according to the manufacturer’s protocol. For each sample, the ratio of firefly to *Renilla* luminescence was measured and normalized to the empty pGL4.24 construct. Two separate biological replicates with at least 3 technical replicates per rs72928038 allele were conducted. For comparison of luminescence conferred by rs72928038 risk and non-risk alleles in the luciferase assay, we used a two-sided Student’s *t*-test.

### Base-editing and PrimeFlow

Base editor mRNA (evoCDAmax-SpCas9-NG) was provided by TriLink Biotechnologies, and was transcribed *in vitro* from PCR product using full substitution of 5-methoxyuridine for uridine. mRNA was capped co-transcriptionally using CleanCap AG analog (TriLink Biotechnologies) resulting in a 5′ Cap 1 structure. *In vitro* transcription reaction was performed as previously described^67^ with the following changes; 16.5 mM magnesium acetate and 4 mM CleanCap AG were used as the final concentration during transcription, and mRNAs were purified using RNeasy kit (QIAgen). Mammalian-optimized UTR sequences (TriLink) and a 120-base polyA tail were included in the transcribed PCR product.

To edit Jurkat T cells, 1×10^6^ were centrifuged at 500 × *g* for 5 min, washed with 1X PBS, and centrifuged again at 500 × *g* for 5 min. The cells were resuspended in 12 uL of plain RPMI 1640, 3 μg of evoCDAmax and 100 μM IDT-synthesized guide RNA was added, and cells were nucleofected using the Neon transfection system program 1600V, 3 pulses, 10ms. Cells were ejected into RPMI. rs72928038 base-edited cells were either left alone or combined with safe harbor base-edited cells (termed WT for the purposes of this study). The cells were incubated for 7 days prior to harvesting for PrimeFlow.

For PrimeFlow, 10 million cells were aliquoted in PBS in polypropylene tubes and centrifuged at 500 × *g* for 5 min. All but 100 μL of the supernatant was discarded (this step is true for every centrifugation step in this protocol) and the cells were resuspended in the residual volume. Cells were then fixed according to the manufacturer’s protocol (ThermoFisher, 88-18005-210) using Fixation Buffer 1 for 30 min rotating at 2–8 °C. Cells were then centrifuged at 800 × *g* for 5 min and the supernatant was discarded. Cells were then permeabilized according to manufacturer’s protocol with addition of RNase inhibitors through inversion, and centrifugation at 800 × *g* for 5 min, then the supernatant was discarded. This step was repeated. A second fixation step was carried out using Fixation Buffer 2 according to manufacturer’s protocol, the samples were mixed, and inverted for 1 h in the dark at RT. The cells were then centrifuged at 800 × *g* for 5 min at RT, and the samples were washed twice with PrimeFlow RNA Wash Buffer, centrifuging the samples at 800 × *g* between each wash for 5 min. The *BACH2* target probe (ThermoFisher) was added at 1X in PrimeFlow RNA Target Probe Diluent, mixed thoroughly by pipetting up and down (100 μL of probe/sample), and incubated at 40 °C for 2 h, with inversion every 30 min. 1 mL of PrimeFlow RNA Wash Buffer was added to each sample, the samples were inverted to mix, and centrifuged at 800 × *g* for 5 min, and the supernatant was aspirated. Samples were then washed with 1 mL PrimeFlow RNA Wash Buffer containing RNase inhibitors twice, followed by centrifugation at 800 × *g* for 5 min. 100 μL of PrimeFlow RNA PreAmp Mix was then added to each sample and briefly vortexed to mix, and the samples were then incubated for 1.5 h at 40 °C with mild vortexing once every 30 min. Samples were washed three times with 1 mL of PrimeFlow RNA Wash Buffer, and then were centrifuged at 800 × *g* for 5 min, and the supernatant was aspirated. 100 μL of PrimeFlow RNA Amp Mix was then added to each sample, the samples were mixed by vortexing, and incubated for 1.5 h at 40 °C with mild vortexing once every 30 min. The cells were then washed twice in 1 mL of PrimeFlow RNA Wash Buffer and centrifuged at 800 × *g* for 5 min. Each sample received 100 μL of PrimeFlow RNA Label Probe diluted in PrimeFlow RNA Label Probe Diluent and incubated for 1 h at 40 °C with mild vortexing once at 30 min. Samples were then washed with 1 mL of PrimeFlow RNA Wash Buffer at RT followed by centrifugation at 800 × *g* for 5 min. The samples were then washed five times with 35 °C PrimeFlow RNA Wash Buffer following each wash with centrifugation at 800 × *g* for 5 min. Samples were then left in 100 μL of PBS and stored in the dark at 4 °C until sorting.

Base edited *BACH2* PrimeFlowed cells were cell sorted into six 10.5% bins, sorting on the extremes of expression (30% on either the low or high portion of the expression distribution, each divided into three contiguous bins each comprising ∼10.5% of the overall distribution) using a Beckman Coulter MoFlo Astrios instrument. For each independent experiment, ∼100K cells were sorted per bin. Genomic DNA for each sample was then reverse-crosslinked using ChIP Lysis Buffer (1% SDS, 0.01 M EDTA, 0.05 M Tris–HCl pH 7.5). Briefly, sorted cells were spun at 800 × *g* for 10 min at 4 °C, the supernatant was aspirated, and the cells were resuspended in 50 μL of ChIP Lysis Buffer, and incubated at 65 °C for 10 min. The samples were then cooled to 37 °C and 2 μL of RNase Cocktail (ThermoFisher, AM2286) was added to each sample and the sample was mixed well by pipetting, followed by incubation at 37 °C for 30 min. 10 μL of Proteinase K (NEB, P8107S) was added to each sample and the sample was mixed well by pipetting, followed by incubation at 37 °C for 2 h and then 95 °C for 20 min. gDNA was extracted using Agencourt XP beads at 0.7X following the manufacturer’s protocol, and the sample was eluted at 100 μL. rs72928038 region amplicon libraries were prepared by nested PCR of each sample, splitting each into two 50 μL reactions (25 μL NEBNext Master Mix, 2.5 μL 10 μM BACH2_gDNAamp_F and 2.5 10 μM BACH2_gDNAamp_R, and 20 μL of reverse-crosslinked sample), program: 98 °C for 30 s, 23 cycles of 98 °C for 15 s, 65 °C for 30 s, 72 °C for 1 min, then 72 °C for 2 min, followed by 1.5X Ampure XP cleanup, and elution in 25 μL EB. A second 50 μL PCR was performed to add Nextera adapters (25 μL NEBNext Master Mix, 2.5 μL 10 μM BACH2_gseq_F and 2.5 10 μM BACH2_gseq_R, and 20 μL of Ampure-cleaned PCR1), program: 98 °C for 30 s, 6 cycles of 98 °C for 15 s, 64 °C for 30 s, 72 °C for 30 s, then 72 °C for 2 min, followed by 1.5X Ampure XP cleanup, and elution in 25 μL EB. A final 50 μL PCR to add Nextera barcodes and sequencing adapters was then performed (25 μL NEBNext Master Mix, 2.5 μL of mixed 10 μM barcoded Nextera_F and Nextera_R, and 22.5 μL Ampure-cleaned PCR2) for 6 cycles; program: 98 °C for 30 s, 25 cycles of 98 °C for 15 s, 62 °C for 15 s, 72 °C for 16 s, then 72 °C for 2 min. The libraries were then quantified by Qubit and TapeStation, mixed at equimolar ratios, PhiX (Illumina, FC-110-3001) was added at 20%, and samples were sequenced aiming to get >1,000,000 reads per bin, on either an Illumina Nextseq 550 or MiSeq.

CRISPResso (version 2.0.29)^68^ was used to count the genotypes of each of the base editor-induced mutations present within the sequencing data associated with each FACS sorting bin. The read counts and genotypes for each sorting bin and the unsorted cells as output by CRIPSResso, were input into R, and MAUDE (version 0.99.3)^69^ was used to infer the expression levels of genotype, separately for each experiment. Here, we assumed that 10.5% of the cells were sorted into each of the sorting bins, which was the approximate number observed to fall into each bin during the experiments. We used MAUDE’s ‘findGuideHitsAllScreens’ function to identify the mean expression associated with each genotype (treating genotypes as MAUDE “guides”), using default parameters. The statistical effect of rs72928038 base edits compared to WT on *BACH2* expression were calculated using a paired (by experiment) one-sided Student’s *t*-test with unequal variance.

### ATAC-seq

We used the FAST-ATAC protocol^11^. Human primary T cells from female subjects (age 12-46) were isolated from blood by Ficoll, followed by flow sorting of live cell single lymphocytes, CD3^+^ CD4^+^. Cells were sorted into RPMI with 10% FBS, and were immediately processed for ATAC-seq. 10,000–40,000 cells were sorted into RPMI 1640 containing 10% fetal bovine serum. The cells were centrifuged at 500 × *g* for 5 min at 4 °C. All of the supernatant was aspirated, ensuring that the pellet was not disturbed in the process. The pellet was then resuspended in the tagmentation reaction mix (25 μL 2X TD Buffer (Illumina, 15027866), 2.5 μL TD Enzyme (Illumina, 15038061), 0.5 μL 1% Digitonin (Promega, G9441), 22 μL H_2_O) and mixed at 300 RPMs at 37 °C for 30 min on an Eppendorf Thermomixer. Immediately after the incubation, samples were purified using a minElute kit (Qiagen, 28006), eluting in 10 μL. The entire sample underwent PCR amplification (a 50 μL reaction with 25 μL NEBNext, 2.5 μL of mixed Nextera_F and Nextera_R (10 μM each; Supplementary Table 14), 10 μL of tagmented DNA, and 12.5 μL H2O) for five cycles with the following program (72 °C, 5 min; 98 °C, 30 s; five cycles of 98 °C, 15 s, 63 °C, 15 s, 72 °C, 1 min). We performed qPCR with 5 μL of the sample to determine the number of additional cycles required, while the rest of the sample remained on ice. The 5 μl of sample was added to a qPCR mix (5 μL of PCR, 5 μl of NEBNext, 0.5 μL mixed Nexter_F and Nextera_R primers, 0.09 μL of 100X SYBR (Invitrogen, S7563), 4.41 μL H2O) and qPCRed (98 °C, 30 s; 20 cycles of 98 °C, 15 s, 63 °C, 15 s, 72 °C, 1 min). The number of cycles that it took to reach 1/3 the maximum fluorescence threshold in the qPCR was then applied via PCR to the original PCR sample. Libraries were cleaned using 1.5X Agencourt XP beads and ethanol washes per manufacturer’s protocol. The DNA concentration of the sample was measured using Qubit and the average fragment size was determined using a TapeStation. Samples were then multiplexed and sequenced using 50 bp paired end chemistry at an average read-count of 30M reads per sample.

Paired-end ATAC-seq reads were mapped to the genome (hg19) using Bowtie2^70^ (v2.2.1; parameters: --maxins 2000), with duplicate reads removed using Picard (v2.20.6; MarkDuplicates REMOVE_DUPLICATES=true), and peaks called using HOMER (v4.6; findPeaks -style dnase).

The comparison of the allelic bias of risk and non-risk alleles in accessible chromatin data from heterozygous individuals was conducted through the use of a binomial test for each sample. Samples indicated in red (Fig. 4C) had significant allelic bias.

### Creation of Bach2 enhancer mutant and knockout mice

All animal procedures were performed in accordance with The Jackson Laboratory, Broad Institute, and Benaroya Research Institute Institutional Animal Care and Use Committees.

The mouse syntenic region including rs72928038 and EH38E2485452 was determined using LiftOver (UCSC) and conservation with human sequence was calculated using EMBOSS Matcher^58^. Bach2^18del^ mice were generated using direct delivery of CRISPR-Cas9 reagents to mouse zygotes following the protocol of Qin et al. including guide design and electroporation^59^. The sequence of guide RNA IDT1038 and ssDO Donor10653 to create Bach2^18del^ is listed in Supplementary Table 14. crRNA containing target sequence (Alt-R CRISPR-Cas9, IDT, 1072532) and tracRNA (Alt-R tracrRNA, IDT,1072534) were hybridized according to the manufacturer’s protocol. The guideRNA duplex was mixed with AltR-SpCas9 V3 (IDT, 108105) in TE (pH 7.5) and incubated for 20 min at room temperature followed by adding ssDO and centrifuging at 14,000 RPM in a microcentrifuge. The final concentrations of the gRNA, spCas9 and ssDO were 600 ng/µl, 500 ng/µl and 1000 ng/µl respectively. 10 µl of the supernatant of the mixture was mixed with 10 µl of Opti-MEM (Thermo Fisher, 3198570) and zygotes treated with acidic Tyrode’s solution (Millapore-Sigma, T1788) for 10 sec and washed with pre-warmed M2 media (Millapore-Sigma, M7167). Electroporation was performed by using ECM830 Square Wave Electroporation System (BTX, 45-0661) with 1 mm electroporation cuvette (Harvard Apparatus, 45-0124) in the following conditions: 2 × 1 ms pulses at 30 V with 100 ms interval. After recovery in 100 µl of M2 media, embryos were transferred into B6Qsi5F1 pseudopregnant female mice.

Genotypes of founder mice were checked by Illumina sequencing. A 384 bp genomic fragment surrounding the target reference (mm10, chr4:32263658-32264041) was amplified from genomic DNA isolated from tail and peripheral blood using 1 µl of prepped DNA in 20 µl of PCR reaction containing 0.4 µl of PrimerStar GXL (TAKARA Bio, R050A), 4 µl of 5× Buffer, 2 µl of 10 mM each dNTP mixture, 0.5 µl each of Bach2ilmn_F1 and Bach2ilmn_R1 primers (10 µM, Supplementary Table 14) with the following conditions: 98 °C for 2 min, 20 cycles of (98 °C for 10 sec, 62 °C for 15 sec, 68 °C for 60 sec), 68 °C for 2 min. PCR products were purified by using 1.1× volume of AMPure XP and eluted by 20 µl of EB. P5/P7 tags were added using 10 µl of first PCR product in a 50 µl of PCR reaction containing 25 µl of Q5 NEBNext master mix, 2.5 µl each of TruSeq multiplexing primers (10 µM) with the following conditions: 98 °C for 30 sec, 6 cycles of (98 °C for 10 sec, 62 °C for 20 sec, 72 °C for 30 sec), 72 °C for 5 min. Amplified products were purified by using 1.2 × volume of AMPure XP, eluted by 20 µl of EB and pooled with the same amount. The mixed library was sequenced on MiSeq (Illumina) using 2 × 250 bp chemistry of nano v2 reagent.

### Isolation of primary mouse T cells

Mouse primary naïve CD8 T cells were isolated from the spleens of WT or Bach2^18del^ mice through sorting on live single lymphocytes, CD3^+^ CD8^+^ CD62L^hi^ CD44^lo^ into PBS containing 2% of FBS. Cells were spun at 500 × *g* for 5 min and lysed by RLT buffer with 40 mM DTT, followed by processing for BRB-seq.

### BRB-seq

Naïve T cells were sorted from spleens collected from 21-week-old females. 5 x 10^5^ cells for each replicate were sorted by using BD FACSymphony S6 with a 70 µm nozzle. The fluorophore-conjugated antibodies and dilutions used for cell sorting are listed in Supplementary Table 14. Total RNA from sorted cells was isolated by using RNeasy plus micro (QIAGEN, 74034). 50 ng for each sample was used for the reverse transcription with barcoded primer BU3 (IDT; Supplementary Table 14) followed by the purification, second strand synthesis and tagmentation following the original BRB-seq protocol but using AMPure XP for purification^34^. Tagmented library was amplified with P5_BRB and BRB_Idx7N5 primers (5 μL, Supplementary Table 14) using NEBNext UltraTM II Q5 Master Mix (NEB, M0544L) which was incubated at 98 °C for 30 sec before adding DNA with the following conditions: 72 °C 3 min, 98 °C for 30sec, and 15 cycles of (98 °C for 10 sec, 63 °C for 30 sec, 72 °C 60 sec), 72 °C for 5 min. Libraries were sequenced by NextSeq 550 High Output with 21 bp for read 1 and 72 bp for read 2 (Illumina). Sequenced reads were aligned using STAR (v2.7.6a, --outFilterMultimapNmax 1)^71^ followed by demultiplexing using BRB-seq Tools (v1.6)^34^.

### BRB-seq differential expression and gene set enrichment analysis

Counted unique UMIs for genes were normalized using variance stabilizing transformation (vst) in DESeq2 (v1.26.0) using default parameters^72^. Differential expression of genes was calculated using DESeq2 with default parameters and genes were then sorted by their differential expression test statistic as input into Gene set enrichment analysis (GSEA).

Expression heatmaps of differentially expressed genes in the Bach2^18del^ mouse compared with WT littermates in Fig. 5e are based on row and column-normalized gene expression. Expression heatmaps of genes from CD8 Tscms with Bach2 gRNA-knockdown vs. empty vector control cells in Supplementary Fig. 11c are similarly based on row and column-normalized gene expression. Comparisons of gene expression for sample genes (Fig. 5d and 5h) were performed by plotting expression levels of each gene from DESeq2 normalized counts from WT and Bach2^18del^ naïve CD8 T cells, followed by performing a Student’s unpaired t-test. Differences in CD62L mean fluorescence intensity between WT and Bach2^18del^ naïve CD8 T cells (Fig. 5e) was performed using an unpaired Student’s *t*-test.

GSEA v4.1.0^73^ was performed from the differential expression ranked gene list. Mouse genes were collapsed to their human orthologs. The GSEAPreranked tool was used with minimum gene set size of 15 and maximum of 250; otherwise, all default parameters were used. The ImmunoSigDB immunologic signatures database (v7.2)^74^ was used for the gene sets in addition to gene sets comprised of differentially expressed genes from the re-analysis of the *Bach2*-perturbed Tscm RNA-seq data (as described below).

### *Bach2*-perturbed Tscm RNA-seq analysis

*Bach2*-gRNA perturbed (n=3) and cognate empty vector control (n=3) RNA-seq data from Tscms^10^ was downloaded from NCBI GEO (GSE152379). Transcript quantification from raw RNA-seq data was performed using Kallisto v0.46.0^75^ against the reference *Mus musculus* transcriptome index based on GRCm38 as provided by the software developers. Quantification of transcripts was performed using parameters --single -l 200 -s 20. Differential expression of genes in *Bach2*-gRNA perturbed vs. empty vector was performed using DESeq2 as described above. Gene sets were constructed from the differential expression results based on genes with a Benjamini-Hochberg adjusted P value < 0.05). One gene set was comprised of the top 200 genes with increased expression in the gRNA vs. empty vector, and the other gene set consisted of the top 200 genes with decreased expression in gRNA vs. empty vector.

### Data visualization and processing

Data visualization, exploratory data analysis, and processing were performed using R v3.6.2.

## Materials Availability

- The Bach218del (stock #35028) mouse strain is available from the Jackson Laboratory (Bar Harbor, ME).
- Plasmids generated in this study will be deposited to Addgene upon publication.

### Data and code availability

Datasets supporting this manuscript are freely available upon reasonable request from the corresponding author, and will be published on NCBI GEO upon the manuscript’s acceptance.

### Code availability

Code supporting this manuscript is freely available upon reasonable request from the corresponding author, and will be published on GitHub upon the manuscript’s acceptance.

## ACKNOWLEDGMENTS

We gratefully acknowledge the contribution of Richard Maser and Genetic Engineering Technologies Service, Jeniffer Kelmenson and Transgenic Genotyping Service, William Schott and Flow Cytometry Service, and Ryan Lynch and Genome Technologies Service at The Jackson Laboratory, Anton McCaffrey and Jordana Henderson at Trilink Biotechnologies, the Broad Institute vivarium, Flow Cytometry Core, and Genomics Core, and the Benaroya Research Institute vivarium and Flow Cytometry Core for expert assistance with the work described in this manuscript. We thank Dr. Peter Gregersen and Dr. Betty Diamond for providing genotyped human PBMCs, and Dr. Susan Malkiel for help with processing human PBMCs for ATAC-seq and for review of the manuscript. We thank Dr. Virginia M. Green for her critical review of the manuscript. This work is funded by U.S. NIH K22 AI153648-01 (J.P.R.), R25 NS065745 (M.H.G.), U01 AI142756 (D.R.L.), RM1 HG009490 (D.R.L.), and R00 HG008179 (R.T.). G.A.N. acknowledges a Helen Hay Whitney postdoctoral fellowship. M.G. is the recipient of an EMBO Long-Term Fellowship (ALTF 486-2018) and a Cancer Research Institute/Bristol-Myers Squibb Fellow (CRI2993).

## AUTHOR CONTRIBUTIONS

J.P.R., R.T., K.M., and M.H.G. conceived the study. J.P.R. performed MPRA, ATAC-seq on human CD4 T cells, base editing experiments, and luciferase, with the help of M.G. K.M. created the Bach2^18del^ mouse line and performed RNA-seq on mouse naïve CD8 T cells. M.H.G, C.G.B., K.M., R.T., J.P.R. performed data analysis. G.A.N. and D.R.L. provided essential base editing reagents and critical advice for base editing experimental design. J.P.R. and M.H.G. wrote the manuscript with the help of K.M., R.T., and N.H. All authors have read and approved the manuscript.

