## Supplementary Information for "Prioritization of autoimmune disease-associated genetic variants that perturb regulatory element activity in T cells"

**Contents:**

Supplemental Figs. 1 - 11

Captions for Supplemental Tables 1 - 14

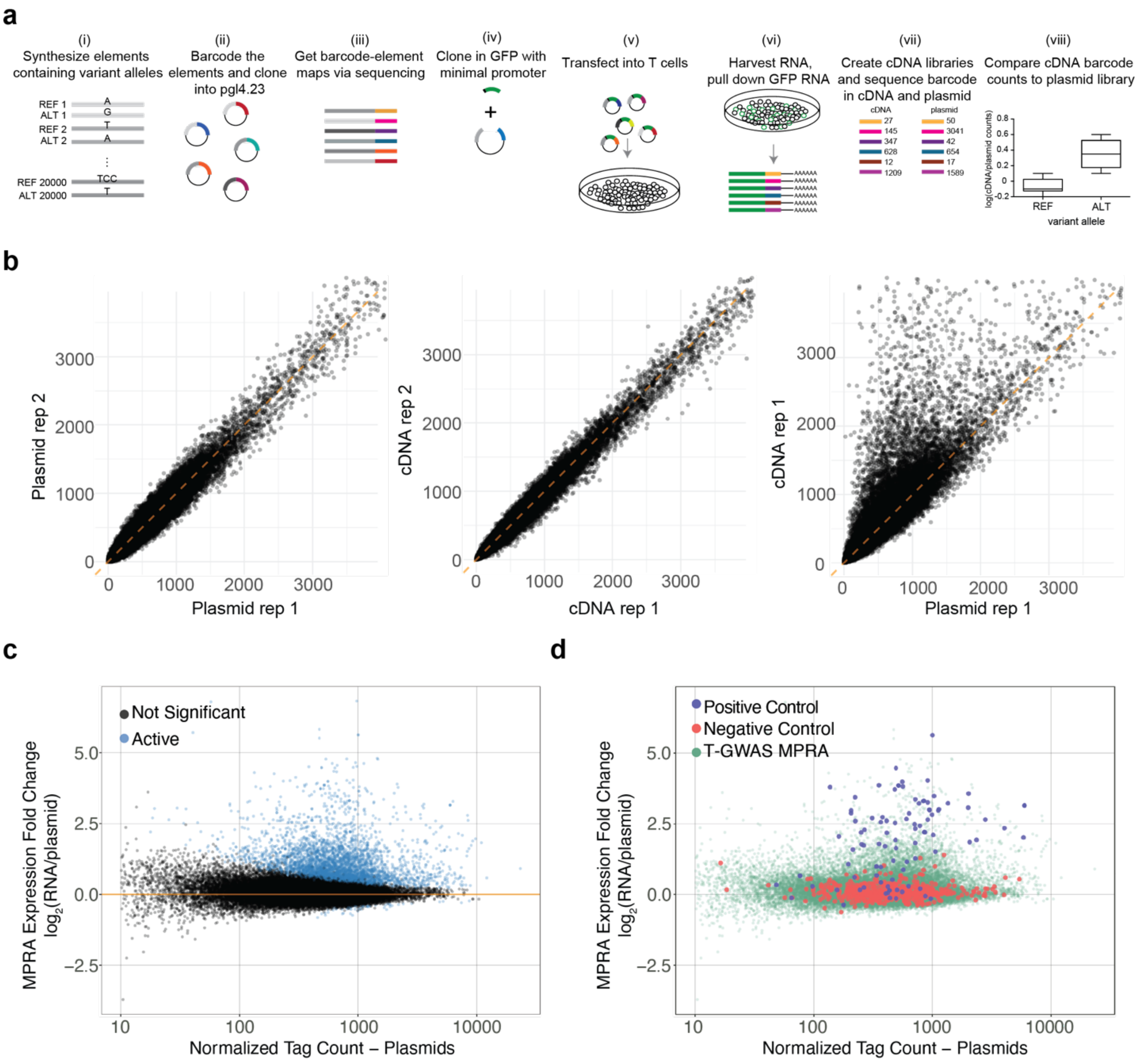


**Supplementary Fig. 1. T-GWAS MPRA workflow and quality metrics. a**) Extended MPRA workflow. i) Oligonucleotide synthesis of elements containing variants and 200 bp surrounding genomic region; ii) barcode elements through PCR; iii) sequence barcoded elements to link barcodes to elements; iv) insert minimal promoter and GFP between element and barcode; v) transfect library into Jurkat T cells; vi) harvest RNA and pull down GFP mRNA; vii) create cDNA and plasmid sequencing libraries and sequence, comparing the prevalence of barcodes in cDNA to their prevalence in plasmid libraries; viii) compare alleles for differential reporter expression. **b**) Correlation of barcode prevalence in separate replicates of plasmid libraries (left), biological replicates of cDNA libraries (middle), and between a cDNA library and a plasmid library (right). **c**) Plot of MPRA expression fold change ($\log_2$ RNA/plasmid; y-axis) against normalized plasmid tag counts for elements with putative cis-regulatory activity (active; blue) and inactive elements (black) within the T-GWAS library. **d**) Plot of MPRA expression fold change ($log_2$ RNA/plasmid; y-axis) against normalized plasmid tag counts for positive controls and negative controls compared to T-GWAS elements. Enrichment calculations for (**b** and **c**) were performed as risk ratios (see Methods), and. P values were calculated based on a two-sided Fisher's exact test. Multiple testing correction for (**b** and **c**) was performed using Bonferroni correction by taking an alpha of 0.05 divided by the number of genomic annotations tested.

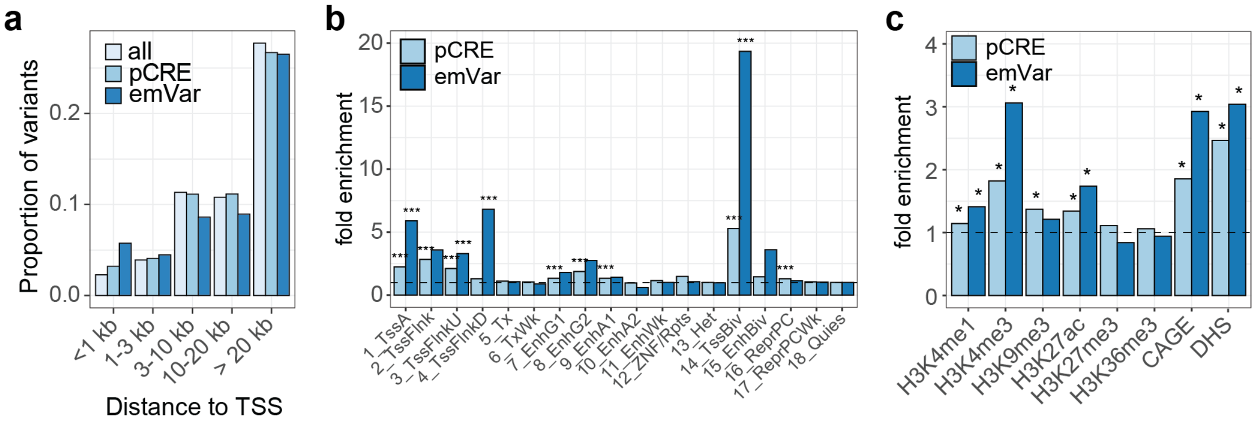


**Supplementary Fig. 2. Variant locations relative to cis-regulatory features. a**) Location relative to TSSs of MPRA tested variants (all), variants on active elements (pCRE), and emVars. **b**) Enrichment of variants within pCREs (light blue) and emVars (dark blue) within chromHMM-defined genomic regions in primary human T cells. * for nominally significant enrichment P value < 0.05; *** for enrichment P value < 1.4 x $10^{-3}$ (0.05 corrected for 36 independent tests). **c**) Enrichment of variants within pCREs and emVars within regions with histone marks (as determined by ChIP-seq), CAGE, and DHS from T cells. *** for enrichment P value < 6.3 x $10^{-3}$ (0.05 corrected for 8 independent tests). Enrichment calculations were performed as risk ratios (see Methods), and P values for (**b** and **c**) were calculated based on a two-sided Fisher's exact test. Multiple testing correction for (**b** and **c**) was performed using Bonferroni correction by taking an alpha of 0.05 divided by the number of genomic annotations tested.

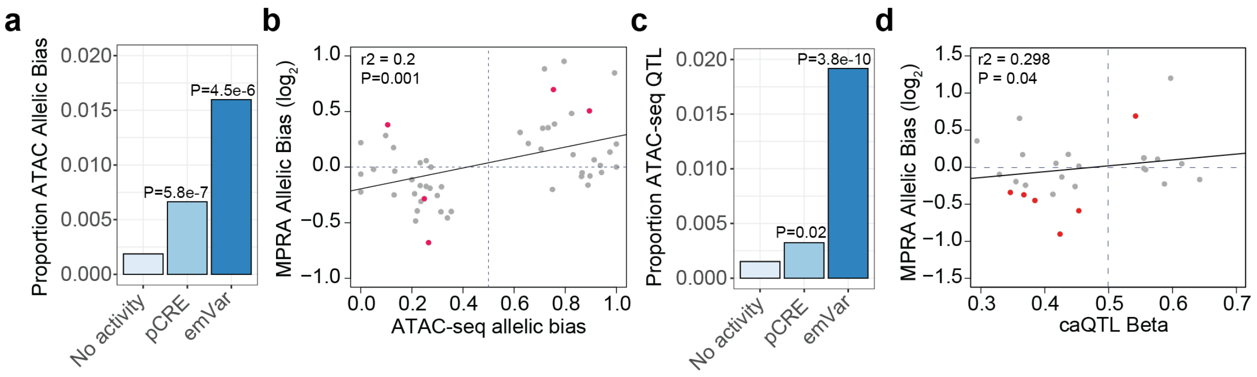


**Supplementary Fig. 3. emVar prevalence for allele-specific chromatin accessibility. a**) Proportion of inactive MPRA variants, MPRA variants within pCREs, and emVars that have allelic bias in ATAC-seq. **b**) Scatter plot comparing MPRA $\log_2$ allelic bias (y-axis) with allelic bias in ATAC-seq from hematopoietic cells (x-axis)[12]. emVars are shown in red dots (n=5) and MPRA variants in pCREs are shown in gray dots (n=45). **c**) Proportion of inactive MPRA variants, variants within pCREs, and emVars that are chromatin accessibility QTLs (caQTLs) from T cells[65]. **d**) Scatter plot comparing caQTL effect size (beta; x-axis) and MPRA $\log_2$ allelic bias (y-axis). emVars are shown in red dots (n=6) and MPRA variants in pCREs are shown in gray dots (n=22).

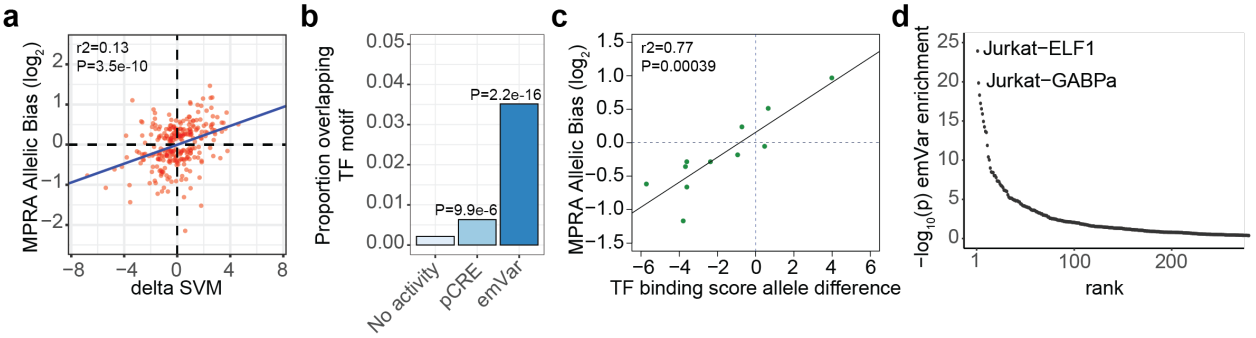


**Supplementary Fig. 4. emVars disrupt motifs that mark active chromatin elements and TFs. a**) Scatter plot comparing delta SVM score (x-axis) with MPRA $log_2$ allelic bias (y-axis) for all single nucleotide substitution emVars (n=278). Pearson's correlation $r^2$ =0.13; correlation P value 3.49 x $10^{-10}$. **b**) Proportion of inactive MPRA variants, variants in pCREs, and emVars that overlap TF motifs. **c**) Scatter plot comparing allele-specific TF binding scores (y-axis) and MPRA $log_2$ allelic bias (x-axis) for emVars predicted to perturb TF binding (n=11). Details of TF motif disruption for all MPRA variants are shown in Table S5. **d**) HOMER enrichment of emVars for overlap with TF ChIP-seq. Y-axis shows -$log_{10}$ enrichment P value (calculated as described in Methods). TF ChIP-seq datasets are ranked from most significant (left) to least significant (right). Full HOMER enrichment results are shown in Table S6.

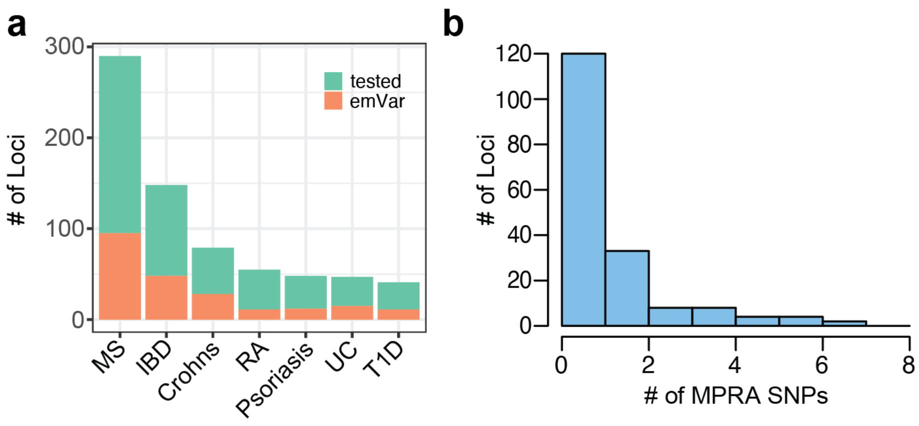


**Supplementary Fig. 5. MPRA prioritizes variants in hundreds of loci. a**) Total number of GWAS loci tested (green) and number of loci with at least one emVar identified (orange) for each disease GWAS. **b**) Histogram of the number of emVars within each GWAS locus.

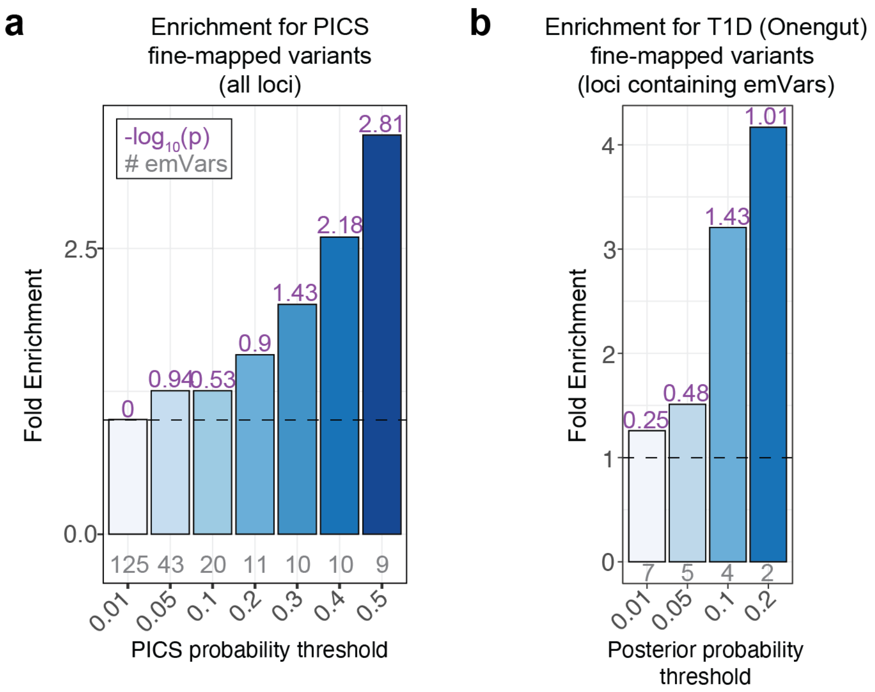


**Supplementary Fig. 6. emVars enrich for high posterior probability fine-mapped variants.** **a**) Bar plot showing enrichment of emVars for PICS statistically fine-mapped variants at all GWAS loci regardless of whether an emVar was detected, with the minimum PICS probability threshold indicated on the x-axis and fold enrichment shown on the y-axis. Details of PICS enrichment results are shown in Table S10. **b**) Bar plot showing enrichment of emVars for fine-mapped T1D GWAS loci from Onengut-Gumuscu et al. Statistical fine-mapping posterior probability threshold is shown on the x-axis and fold enrichment shown on the y-axis. For both **a**) and **b**), gray numbers below each bar show the number of emVars that are statistically fine-mapped at a given PICS probability threshold. Purple numbers above each bar show the $-\log_{10}$ of the enrichment P value. Enrichment in (**a**) and **(b)** were calculated as a risk ratio (see Methods), and P values were determined through a two-sided Fisher's exact test.

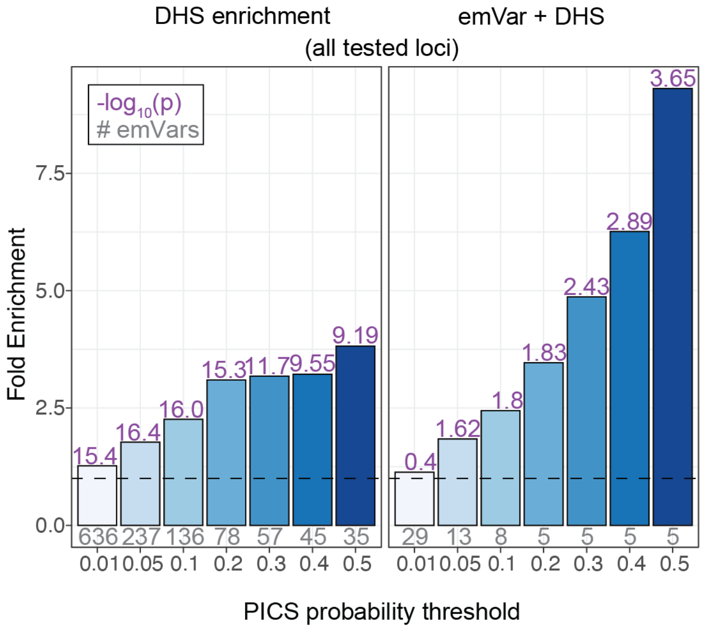


**Supplementary Fig. 7. emVars in accessible chromatin enrich highly for fine-mapped variants.** Bar plot showing enrichment (from all loci tested) of DHS alone (left) and emVars in T cell DHS (right) for PICS fine-mapped variants, with the minimum posterior probability threshold indicated on the x-axis. Gray numbers below each bar show the number of emVars that are statistically fine-mapped at a given PICS probability threshold. Purple numbers above each bar show the $-\log_{10}$ of the enrichment P value. Details of PICS enrichment results are shown in Table S10. Enrichment was calculated as a risk ratio (see Methods), and P values were determined through a two-sided Fisher's exact test.

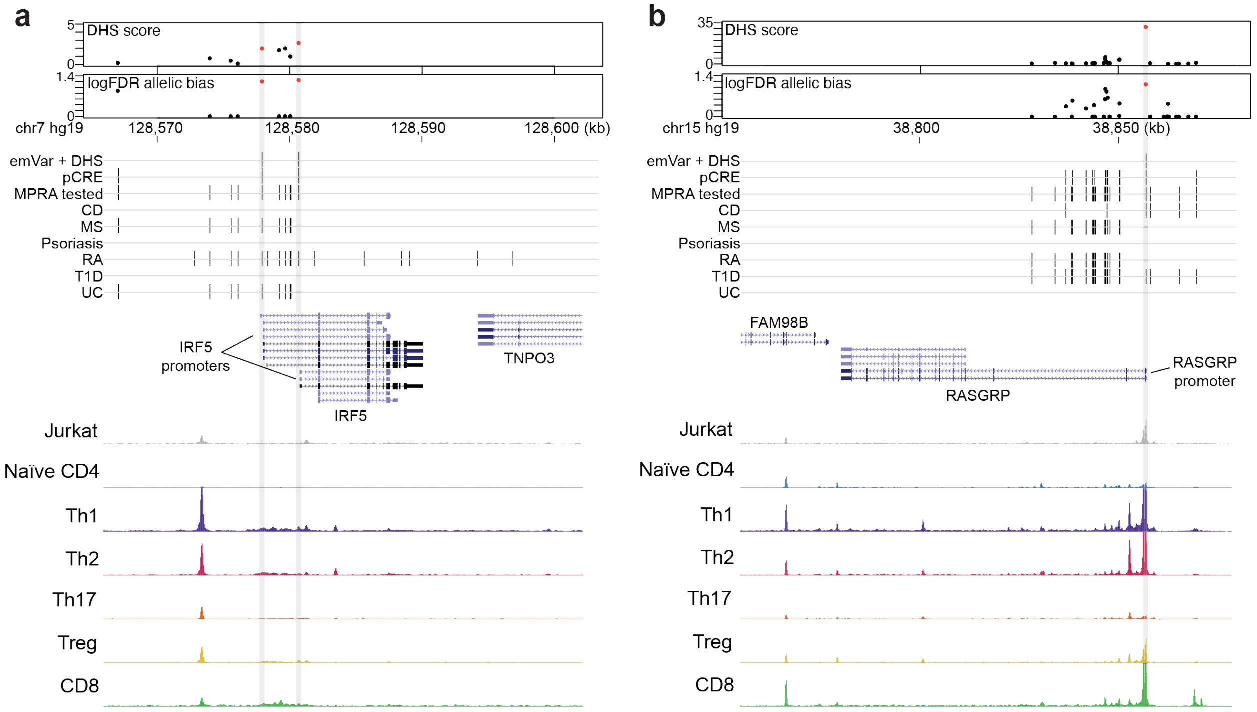


**Supplementary Fig. 8. Putative causal variants in the promoters of *IRF5* and *RASGRP*. a** and **b**) Dotplots showing DHS signal (DHS score) and statistical significance of allelic bias ($log_{10}$FDR of MPRA allelic bias) for MPRA variants in the region; all tested variants on haplotype (black), significant emVars in DHS (red) (top). Position of variants that are emVars, pCREs, variants tested in MPRA, and disease associated variants for CD, MS, psoriasis, RA, T1D, and UC from the GWAS Catalog[56] (middle). Genes in the locus are shown along with chromatin accessibility profiles (from in Jurkat and specific T cell subsets) and T cell pcHiC loops anchored on the region containing the emVar. Gray line depicts position of the prioritized emVar position with respect to all data types. Statistical significance of allelic biases in (**a**) and **(b)** were calculated using a paired Student's two-sided *t*-test as described in Methods.

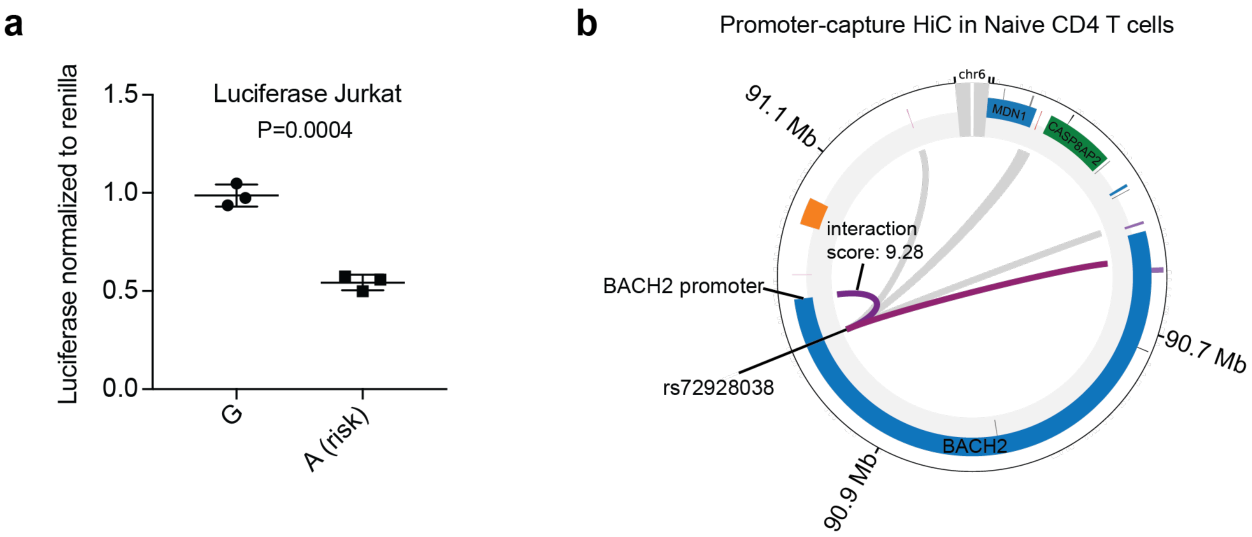


**Supplementary Fig. 9. rs72928038 reduces luciferase reporter expression and contacts the *BACH2* promoter. a**) Luciferase reporter activity of rs72928038 alleles (n=3, two independent experiments). Central tendency is shown as median and all points are plotted to show dispersion. **b**) Promoter capture HiC (pcHiC[23]) conducted in naïve T cells anchored on the region containing the rs72928038. For **(a),** statistical significance was calculated using a Student's two-sided *t*-test.

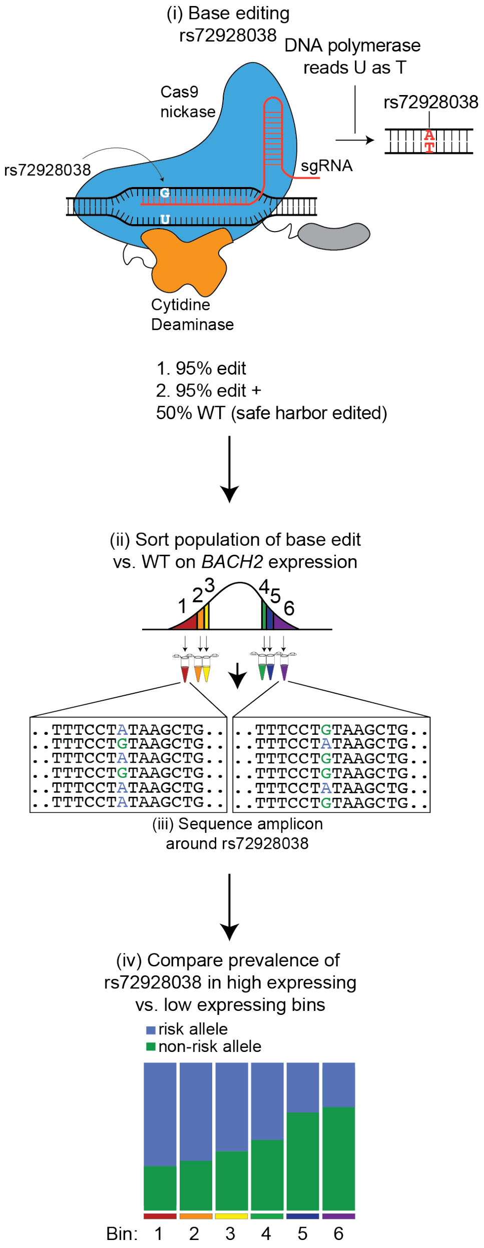


**Supplementary Fig. 10. Base editing of rs72928038 and *BACH2* PrimeFlow.** i) Schematic of installing rs72928038 using the evoCDAmax cytosine base editor, achieving 95% base editing. We also created a second condition, separately combining the 95% base edited cells with WT base-edited cells (combined 50/50) post-nucleofection. (ii) We performed PrimeFlow, staining *BACH2* mRNA, and sorted cells based on high and low *BACH2* expression. iii) We sequenced the amplicon containing rs72928038 in all sorted populations. iv) Mock data of expected ratios of risk vs. non-risk alleles in high and low bins of *BACH2* expression. If rs72928038 reduces *BACH2* expression, one would anticipate the edited risk allele to enrich in low *BACH2* expression bins.

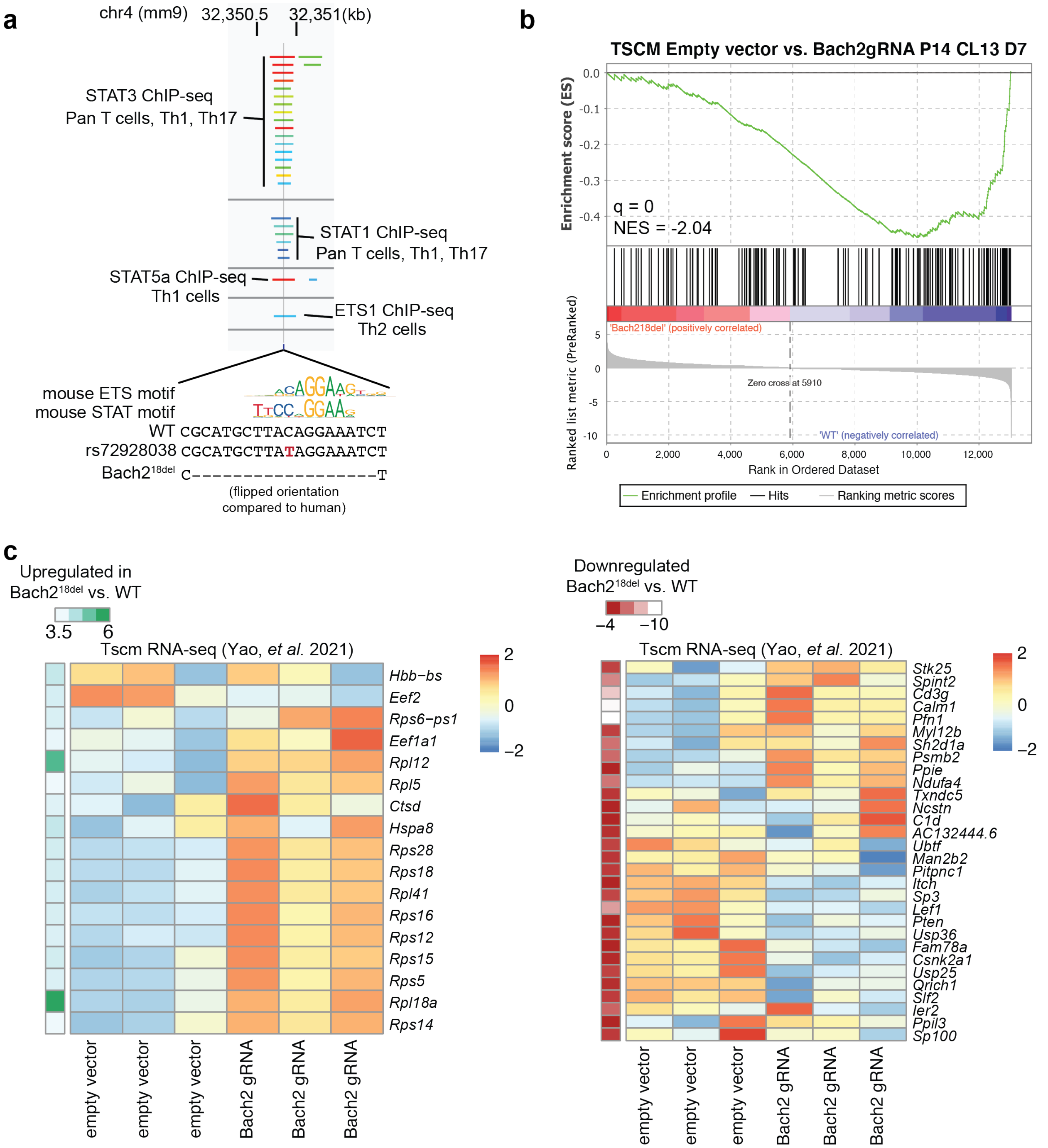


**Supplementary Fig. 11. Orthologous rs72928038 region binds STATs and ETS1 and partially recapitulates transcriptional phenotypes of *Bach2*-deficient Tscms. a**) STAT and ETS1 TF ChIP-seq peaks[33] overlapping mouse rs72928038 ortholog. **b**) GSEA enrichment of Bach2$^{18del}$ vs. WT naïve CD8 T cells. Depicted GSEA results for a gene set derived from genes upregulated in empty vector vs. Bach2 sgRNA-transduced Tscms. Full GSEA results are shown in Table S13. **c**) Expression of genes in Tscms that have been transduced with empty vector or a Bach2 sgRNA (same experiment as in **b**) for differentially expressed genes in Bach2$^{18del}$ vs. WT naïve CD8 T cells. Genes upregulated in $Bach2^{18del}$ T cells as compared to WT are on the left and downregulated are on the right. Normalized enrichment score (NES) in **(b)** was calculated based on observed enrichment as compared to enrichments from permuted data as previously described and statistical significance shown as the false discovery rate (q).

**Supplementary Table 1. GWAS loci analyzed in this study.** MS, T1D, RA, IBD, and psoriasis loci analyzed.

**Supplementary Table 2. MPRA oligos.** 200 bp oligos synthesized with 170 bp genomic sequence containing the variant and 15 bp Gibson Assembly adapters.

**Supplementary Table 3. MPRA results.** All MPRA expression and allelic bias (skew) results.

**Supplementary Table 4. MPRA functional annotation.** T cell MPRA effect, DHS, ATAC allelic skew, caQTL, T cell eQTL, acQTL, and TF motifbreakR information for each variant.

**Supplementary Table 5. MPRA TF motif disruption.** Full details of all MPRA variants with significant predicted disruption of TF position-weighted matrices based on motifbreakR.

**Supplementary Table 6. MPRA HOMER results.** Enrichment of emVar-containing regions for TF ChIP-seq based on HOMER.

**Supplementary Table 7. PICS statistical fine-mapping data.** Full PICS fine-mapping results across all GWAS loci tested.

**Supplementary Table 8. MPRA results with PICS probabilities and credible sets.** Full PICS results by MPRA SNP, including PICS results for strongest GWAS association.

**Supplementary Table 9. emVar PICS enrichment (all loci).** emVar PICS enrichments for all loci, regardless of whether emVar was detected.

**Supplementary Table 10. emVar PICS enrichment (loci containing an emVar):** emVar PICS enrichments for loci containing an emVar.

**Supplementary Table 11. rs72928038 allele-specific ATAC-seq from heterozygous individuals**

**Supplementary Table 12. Bach2$^{18del}$ vs. WT DESeq2 RNA-seq results.** DESeq2 differential expression results for Bach2$^{18del}$ vs. WT mouse naïve CD8 T cells

**Supplementary Table 13. Bach2$^{18del}$ vs. WT gene set enrichment analysis.** GSEA results for differentially expressed genes between Bach2$^{18del}$ vs. WT mouse naïve CD8 T cells

**Supplementary Table 14. Primers and antibodies used in this study**
